# Two synthetic 18-way outcrossed populations of diploid budding yeast with utility for complex trait dissection

**DOI:** 10.1101/2020.01.09.900480

**Authors:** Robert A. Linder, Arundhati Majumder, Mahul Chakraborty, Anthony Long

## Abstract

Advanced generation multi-parent populations (MPPs) are a valuable tool for dissecting complex traits, having more power than GWAS to detect rare variants, and higher resolution than F_2_ linkage mapping. To extend the advantages of MPPs in budding yeast, we describe the creation and characterization of two outbred MPPs derived from eighteen genetically diverse founding strains. We carried out *de novo* assemblies of the genomes of the eighteen founder strains, such that virtually all variation segregating between these strains is known and represent those assemblies as Santa Cruz Genome Browser tracks. We discover complex patterns of structural variation segregating amongst the founders, including a large deletion within the vacuolar ATPase *VMA1*, several different deletions within the osmosensor *MSB2*, a series of deletions and insertions at *PRM7* and the adjacent *BSC1*, as well as copy number variation at the dehydrogenase *ALD2*. Resequenced haploid recombinant clones from the two MPPs have a median unrecombined block size of 66kb, demonstrating the population are highly recombined. We pool sequenced the two MPPs to 3270X and 2226X coverage and demonstrate that we can accurately estimate local haplotype frequencies using pooled data. We further down-sampled the poolseq data to ~20-40X and show that local haplotype frequency estimates remain accurate, with median error rate 0.8% and 0.6% at 20X and 40X, respectively. Haplotypes frequencies are estimated much more accurately than SNP frequencies obtained directly from the same data. Deep sequencing of the two populations revealed that ten or more founders are present at a detectable frequency for over 98% of the genome, validating the utility of this resource for the exploration of the role of standing variation in the architecture of complex traits.

## Introduction

A complete understanding of the genetic basis of complex traits is a goal shared by many disciplines. Although much progress has been made in dissecting the genetic architecture of complex traits such as adaptation, disease susceptibility, human height, and crop performance, a major fraction of standing variation for most traits has remained recalcitrant to dissection (Manolio *et al*. 2009). This is often referred to as the ‘missing’ heritability problem. Rapid progress in addressing the missing heritability problem seems most likely in model systems that can be genetically and experimentally manipulated in a controlled setting. In contrast to humans, in model genetic systems variants of subtle effect can be validated via allele replacement experiments.

One of the mainstays of modern genetic mapping studies has been the use of pairwise crosses between genetically diverged founder strains. Large segregating populations can then be used to map phenotype to genotype. This approach, laid out in its modern form for complex traits was initially described by (Lander and Botstein 1989) and reviewed in (Flint and Mott 2001; Mackay 2001; Liti and Louis 2012) has proven to be especially fruitful in budding yeast, such that mapped QTL tend to explain > 70% of the narrow-sense heritability of most traits (Ehrenreich *et al*. 2010; Bloom *et al*. 2013, 2015, 2019; Märtens *et al*. 2016). However, QTL mapping has suffered both from a lack of resolution and a severe under-sampling of the functional variation potentially segregating in natural populations. In this regard, association studies enjoy much finer mapping resolution and sample a larger proportion of the variation present in a natural population (WTCCC 2007; Visscher 2008). However, large-scale association studies are often under-powered to detect rare alleles (Spencer *et al*. 2009), regions that harbor multiple causal sites in weak LD with one another (Pritchard 2001; Thornton *et al*. 2013), rare or poorly tagged structural variants (Hehir-Kwa *et al*. 2016) or variants that are poorly tagged more generally. Furthermore, as GWAS studies grow to include tens of thousands of individuals they can suffer from false positives from population stratification (Berg *et al*. 2019) or other experimental block artifacts associated with large scale projects (Sebastiani *et al*. 2011; Chen *et al*. 2017).

Advanced generation multiparent populations (MPPs) consisting of recombinants derived from several founder individuals have been proposed as a bridge between pairwise linkage mapping and association studies in outbred populations (“The Collaborative Cross, a community resource for the genetic analysis of complex traits” 2004; Macdonald and Long 2007). MPPs are created by crossing several (inbred or isogenic) founder strains to one another in order to maximize diversity, and then intercrossing the resulting population for several additional generations to increase the number of recombination events in the population. In many model systems Recombinant Inbred Lines (RILs) are derived from the MPP via inbreeding. The resulting homozygous RILs are fine-grained mosaics of the original founding strains that have been successfully used to dissect complex traits in *Arabidopsis thaliana* (Kover *et al*. 2009; Huang *et al*. 2011), *Drosophila melanogaster* (Macdonald and Long 2007; King *et al*. 2012a, 2012b), *Mus musculus* (Aylor *et al*. 2011; Threadgill and Churchill 2012), *Saccharomyces cerevisiae* (Cubillos *et al*. 2013), *Zea mays* (McMullen *et al*. 2009), *Caenorhabditis elegans* (Noble *et al*. 2019), and several other systems (de Koning and McIntyre 2017). MPP RILs are a powerful resource for dissecting complex traits due to increased mapping resolution relative to F2 populations and increased natural variation sampled by the founders. Furthermore, unlike association studies, both rare alleles of large effect segregating among the founders as well as allelic heterogeneity can be detected in MPP RILs (Long *et al*. 2014). Although the majority of studies to date have studied RILs derived from MPPs, it is possible to dispense with the creation and maintenance of RILs and sample the MPP directly (Mott *et al*. 2000; Macdonald and Long 2007), and indeed early MPP efforts did not employ RILs.

Despite the clear advantages of MPPs, only a single MPP has been described in budding yeast (Cubillos *et al*. 2013), which is surprising as this species is ideally suited in many other ways for the dissection of complex traits. Large population sizes can be maintained in a controlled environment and a few rounds of meiosis results in recombination events spaced at near genic resolution. The potential of MPPs in budding yeast was demonstrated by Cubillos et al., who crossed four genetically highly diverged strains and intercrossed the resulting population for twelve generations to generate a highly recombined population that has been shown to be capable of mapping complex traits to high resolution (Cubillos *et al*. 2013, 2017). In order to expand the potential of budding yeast to contribute to our understanding of complex traits we have developed two large, highly outbred populations of budding yeast derived from a cross of 18 genetically diverged founders. Like previous work, populations were intercrossed for 12 generations to produce highly recombined mosaic populations that capture a large amount of the standing variation present in *S. cerevisiae*. Here we describe the derivation of the founders that allows the 18-way cross to be carried out, *de novo* PacBio assemblies of each founder such that all variation segregating in the population is known, and the characterization of ten haploid recombinant clones from each population to estimate the size distribution of haplotype blocks in the MPPs. We further carry out deep short read resequencing of the MPPs, estimate founder haplotype frequencies as a function of location in the genome, and show that at Illumina sequencing coverages as low as ~20X-40X haplotype frequencies can be accurately estimated. The MPPs and tools we derive have great utility for dissecting complex traits in yeast.

## Materials and Methods

### Strains and media

All yeast strains used in this study came from heterothallic, haploid derivatives of a subset of the SGRP yeast strain collection kindly provided by Gianni Liti (Cubillos *et al*. 2009). A list of strains used, relevant genotypes (before and after our modifications), and their geographical origins is shown in Table 1. Additionally, two mate-type testing yeast strains were used (kindly provided by Ian Ehrenreich) that are selectively killed by the presence of either *Mat a* or *Mat α* haploids, but not by diploids. For propagating plasmids, *Escherichia coli* strain DH5α was used according to the manufacturer’s recommendations (Invitrogen). Bacterial transformants were selected on LB agar, supplemented with 100ug/mL ampicillin (‘LBamp’) (Fisher). Nonselective media for growth and maintenance of all yeast strains included rich media consisting of 1% yeast extract, 2% peptone, and 2% dextrose (‘YPD’) (Fisher). For solid media, 2% agar was added. Additionally, media consisting of 1% yeast extract, 2% peptone, 2% glycerol and 2.5% ethanol (‘YPEG’) was used to prevent the growth of *petite* mutants. For selecting yeast transformants, when *Ura3MX* was the marker, synthetic complete drop-out uracil (Sc -Ura) plates were used (Sunrise Scientific). When *KanMX*, *HphMX*, or *NatMX* were the markers used, transformants were selected on YPD plates supplemented with 200ug/mL of G418, 300ug/mL of Hygromycin B (‘hyg’), or 100ug/mL nourseothricin sulfate (‘cloNAT’), respectively. For counterselection of yeast that lost the *Ura3MX* marker, synthetic complete media supplemented with 1mg/mL 5-FOA was used (‘5-FOA’). Two types of sporulation media were used in this study. Type 1 consisted of 1% potassium acetate, 0.1% yeast extract, and 0.05% dextrose (‘PYD’) to which ampicillin was added to a final concentration of 50ug/mL, while type 2 consisted of 1% potassium acetate and a 1X dilution of a 10X amino acid stock (composed of 3.7g of CSM -lysine (Sunrise Scientific) supplemented w/10mL of 10mg/mL lysine in 1L total volume), pH adjusted to 7 (‘PA7’). Just before use, ampicillin was added to PA7 to a final concentration of 100ug/mL.

**Table 1.**
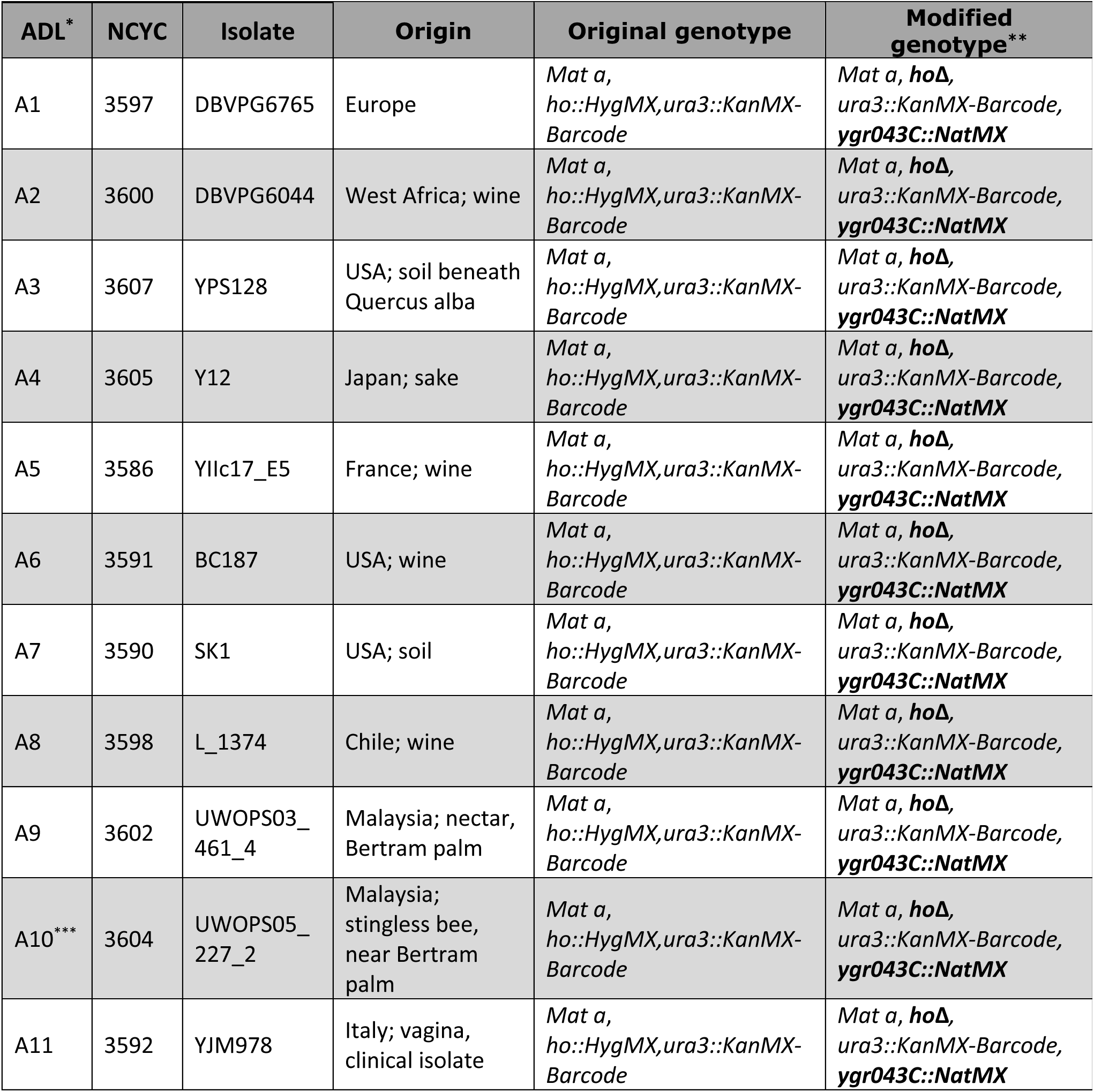
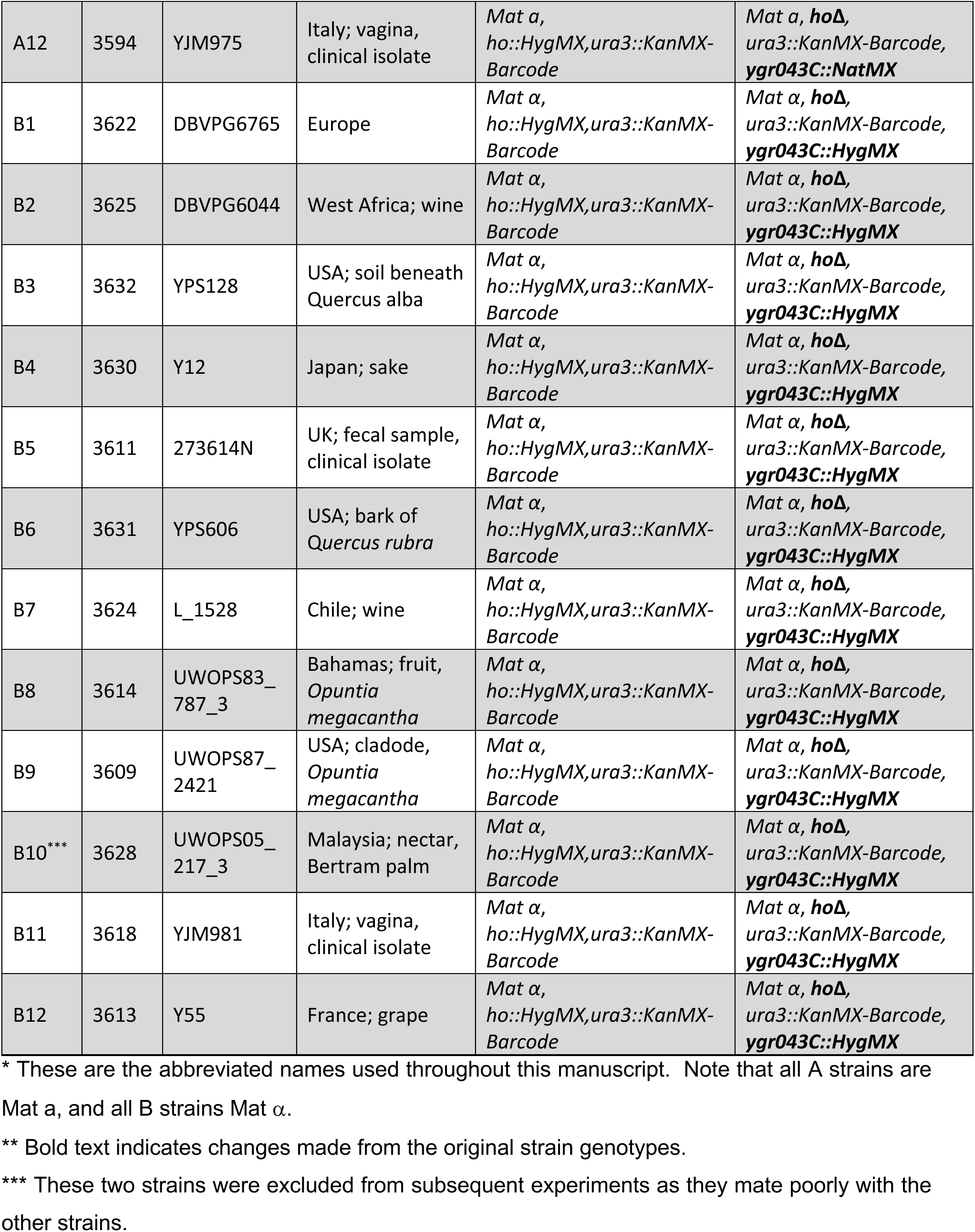
An overview of the strains used in this study.

### Modification of 24 haploid budding yeast strains to create founders for the synthetic population

The strains used in this study were modified by generating clean deletions of the *HO* gene to recover the *HphMX* marker, followed by replacement of a pseudogene, *YCR043C*, which is closely linked to the mating type locus, with either a *NatMX* cassette in *Mat a* haploids or a *HphMX* cassette in *Mat α* haploids. This manipulation was carried out to enable high-throughput selection of diploids.

The *HphMX* marker in *HO* was recovered via transformation with a URA3 cassette flanked with direct repeats and selection on URA-plates followed by selection on 5-FOA plates to recover URA3. The URA3 cassette was assembled from four fragments: a pBluescript II KS(+) backbone linearized with EcoRV and gel purified (for propagation in *Escherichia coli*), the *URA3* gene from *Candida albicans* with flanking 500bp direct repeats from *Aschbya gossypii* (pAG61, addgene #35129), and a 450bp region directly upstream of the *HO* gene, and a 390bp region directly downstream of the *HO* gene. Primers pAG61_HO-F/R were used to amplify *URA3* and the flanking direct repeats, while primers HO-US F/R and HO-DS F/R were used to amplify the regions flanking the *HO* gene from strain DBVPG6765. Primers used in this study are listed in Table S1 and included overhangs to allow for HiFi assembly. The four fragments were assembled using the NEB HiFi Assembly Master Mix according to the manufacturer’s recommendations (NEB), transformed into chemically competent DH5α (Invitrogen), and recovered on LB Amp plates. Recovered URA plasmid cassettes with *HO* flanking sequences were PCR amplified from the plasmid template using primers HO-US-F and HO-DS-R, transformed into all 24 haploid strains using a standard lithium acetate protocol, and plated onto Sc - Ura plates. Single colonies were re-streaked onto Sc -Ura (X2), final colonies were tested for the presence of the *KanMX4* marker and absence of *HphMX4* marker via *G418* and *hyg* plating respectively. O/N cultures of successfully knocked-out transformants were spread onto 5-FOA plates and grown for 2d at 30°C in order to select for cells that had ‘popped out’ the *Ura3* cassette. Single colonies were re-streaked onto 5-FOA plates (2X). DNA was extracted (adapted from CSH handbook, p. 116) from the resulting colonies and DNA amplicons spanning the *HO* locus were obtained and Sanger sequenced to confirm the clean deletion of the *HO* gene.

To delete *YGR043C* in the 24 newly generated haploid *hoΔ Ura3::KanMX4* strains, oligos were ordered from IDT that amplify the entire MX4 cassette, including the promoter and terminator regions, and were tailed with 100 bases of homology to the regions immediately upstream and downstream of the *YGR043C* CDS. Either pAG32 (addgene #35122) or pAG25 (addgene #35121) were used as a template to generate knock-out constructs that incorporate the *HphMX4* or *NatMX4* cassettes, respectively. PCR reactions were cleaned up to remove unamplified circular plasmid template by gel extraction followed by digestion with DpnI and a PCR cleanup reaction (Qiagen PCR Purification kit). *Mat a* yeast were then transformed with the *cloNAT* resistance cassette, while *Mat α* yeast were transformed with the *hyg* resistance cassette using the standard lithium acetate protocol and selecting on YPD supplemented with cloNAT and G418 or hyg and G418, respectively. This double selection with G418 was done to ensure that cassette swapping had not occurred. To ensure *YGR043C* had been correctly replaced in each strain, the region was amplified and Sanger sequenced. All 24 newly generated strains were checked again for *HO* deletion using the HO-big-flank-F/R primers. The strains were also checked to ensure they had maintained the correct barcodes originally inserted in Cubillos et al throughout all the manipulation steps by amplifying the barcodes using the barcode-check-F/R primer pair and Sanger sequencing the amplicons using the M13(-47)F primer. As a final check, all 24 haploid strains were streaked onto YPD supplemented with hyg and cloNAT to ensure that none of the strains could grow on both antibiotics.

### 18-way crossing scheme, version 1

A full diallele cross of eleven *Mat a* and eleven *Mat α* strains (excluding strains A10 and B10) was carried out (with four strains in common). A schematic of the mating scheme is shown in Figure 1A, while Table 1 lists the strains used in this study. Strains A1-A5 and A6-A12 (excluding A10) were struck in horizontal rows onto two YPD plates each (total of 4 YPD plates), then strains B1-B5 and B6-B12 (excluding B10) were each struck in vertical rows onto two of the YPD plates such that each ‘B’ strains intersected with each ‘A’ strains. All 121 pairwise combinations of the ‘A’ and ‘B’ strains were thus represented across the four YPD mating plates. Mating occurred O/N at 30°C after which diploids were selected by replica plating onto YPD plates with hyg and cloNAT. A single colony from each of the 121 crosses was then incubated O/N in YPD with hyg and cloNAT at 30°C at 180RPM. An equal volume of each culture and 30% glycerol was used to make frozen stock that was then archived at −80°C. An equal volume from each diploid culture was then combined to make the 18-way population, which was washed twice with PYD + amp then split into two 1L flasks with 200mL total PYD + amp each. Sporulation was carried out for 5d *en mass* at 30°C at 180RPM to complete the first round of out-crossing.

**Figure 1.**
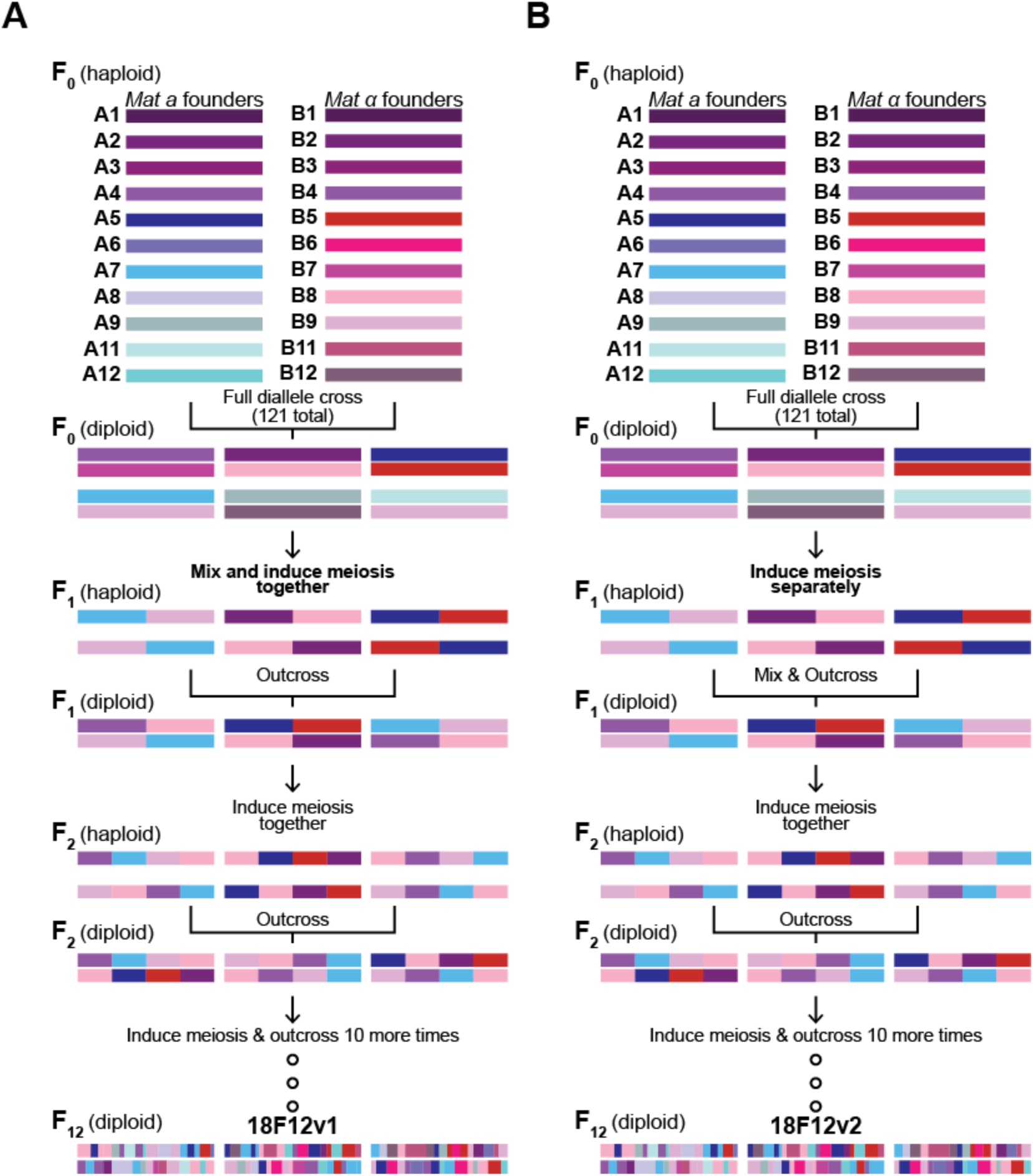
Schematic of the outcrossing process used to make the two 18F12 diploid populations. Both populations were established by a full diallele cross of all 22 isogenic haploid founder strains. A1/B1, A2/B2, A3/B3, and A4/B4 are different mating types of the same strains and are the same strains used in (Cubillos *et al*. 2013). In (**A**), all pairwise crosses were mixed before the first round of sporulation. This is in contrast to (**B**), in which mixing did not occur until after an initial sporulation event. In both cases, mixed populations were taken through additional rounds of sporulation and random mating for a total of 12 meiotic generations.

### Additional outcrossing in the 18-way cross, version 1

Eleven additional cycles of mass sporulation followed by random mating were carried out for a total of twelve rounds of outcrossing for both replicates (Table S3). After sporulation, 50mL of culture was spun down at 2000g for 2m and resuspended in 1mL of Y-PER. Samples were transferred to a 1.5mL centrifuge tube and vortexed. Cells were washed twice and resuspended in 500uL of ddH_2_O with 5uL of 5U/uL zymolyase. The tubes were shaken vigorously in the Geno Grinder 2000 at 750 shakes/minute for 45m. 500uL of 400um silica beads were then added to the samples, which were again put in the Geno Grinder for 5m at 1500 shakes/minute. The supernatant was transferred to a fresh 1.5mL centrifuge tube, washed once in YPD, resuspended in 500uL of YPD, and transferred into 50mL of YPD in a 1L flask. Mating was carried out O/N at 30°C at 40RPM. The next day, mated cells were harvested, transferred to YPD with cloNAT, hyg, and amp, and incubated O/N at 30°C at 180RPM. The next day, 7mL from the O/N culture was used to make glycerol stock while 5mL was harvested, washed twice, and resuspended in 200mL of PYD + amp. Sporulation was carried out for 5d at 30°C at 180RPM (see Table S2). If the experiment had to be paused, 5mL of glycerol stock from the most recently completed cycle was used to begin the next cycle of sporulation.

### 18-way crossing scheme, version 2

A full diallele cross of the same eleven *Mat a* and eleven *Mat α* founder strains used to create 18F12v1 was again carried out. To initiate this process, an equal volume from cultures containing each *Mat a* founder strain were mixed with each *Mat α* founder strain in all 121 possible pairwise combinations in 24-well deep-well plates (hereafter ‘24DWP’) in a total volume of 1mL of YPD (no ampicillin added) (see Note S1). Mating was carried out in liquid culture for 4-5h at 30°C at 50RPM, after which mating was verified by checking for the presence of zygotes and/or shmooing under a microscope. At this point, 1mL of YPD supplemented with 200ug/mL cloNAT and 600ug/mL hyg was added to each culture (the final concentrations of cloNAT and hyg were 100ug/mL and 300ug/mL, respectively) to select for successfully mated diploids and incubated O/N at 200RPM at 30°C. After O/N selection, 140uL from each culture was combined w/140uL of 30% glycerol to make frozen stock of each cross. The remaining cultures were harvested at 3000 RPM for 5m and the pellets were washed then resuspended in 4mL of PA7 + amp. Sporulation was carried out for 6d at 30°C at 275 RPM in the 24DWPs.

All 121 sporulating cultures were checked using a microscope to determine the amount of sporulation that occurred; cultures were graded on a scale of 0-5 with 0 being no sporulation and 5 being almost complete sporulation. Crosses that did not sporulate were excluded from subsequent steps (see Table S2). After checking for sporulation, cultures were harvested, washed, then resuspended in 500uL of spore isolation solution (hereafter ‘SIS’: 25U zymolyase + 10mM DTT + 50mM EDTA + 100mM Tris-HCl, pH 7.2 up to 500uL) and incubated for 1h at 30°C at 250 RPM to spheroplast cells. Cultures were then harvested and resuspended in 1% Tween 20 to selectively lyse unsporulated cells. Following this, cultures were again harvested and resuspended in 500uL of spore dispersal solution (hereafter ‘SDS’: 1mg lysozyme + 5U zymolyase + 1% Triton X-100 + 2% dextrose + 100mM PBS, pH 7.2 up to 500uL). Cultures were transferred to Eppendorf tubes with 500uL of 400um beads and bead milled using a Geno Grinder 2000 at 1500 strokes per minute for 5m to break up tetrads after which all cultures were placed at 4°C O/N. The next day, all tubes were vortexed at high speed for 30s, the supernatant was transferred to a 24DWP, 500uL of 100mM PBS, pH 7.2 was added back to the beads, followed by vortexing for 30s and transferring to the same wells of a 24DWP to maximize recovery of spores from the beads. Cultures were washed once in PBS, then resuspended in 500uL of 100mM PBS, pH 7.2 and 100uL was transferred to a 96-well clear plate to measure the OD630 of each culture in duplicate using a BioTek Synergy HT plate reader. OD630 measurements were then used to normalize the density of spores from each cross that were pooled together (see Note S2). The spore pool was washed twice with 5mL of YPD, then resuspended in 12.5mL of YPD. This culture was split in half and transferred to two 250mL flasks, each with 6.25mL of YPD, to establish two replicate populations. Mating was carried out O/N at 30°C with gentle shaking at 40 RPM. The next day, 12.5mL of YPDach (with a 2x mix of ampicillin, cloNAT, and hyg) was added to each culture to select for diploids. Cultures were incubated O/N at 200 RPM at 30°C. This established replicate F_2_ populations of the 18-way cross, version 2 (hereafter ‘18F2v2’). The following day, 7mL of the replicate populations were frozen down at −80°C with an equal volume of 30% glycerol. The remaining volume was spun down and used to initiate a second round of outcrossing.

### Additional outcrossing in the 18-way cross, version 2

Eleven additional cycles of mass sporulation followed by random mating were carried out for a total of twelve rounds of outcrossing for both replicates. As replicate 2 was treated differently during a couple of cycles, replicate 1 was the population chosen for subsequent analyses and, as such, will be the only replicate of version 2 described further. Each cycle consisted of 3-6d of sporulation after which diploids were randomly mated for 3-4h. This was followed by an O/N selection step in YPDach to enrich for mated diploids. After selection, an aliquot of each population was frozen down at −80°C with the remaining culture used to initiate the next cycle of outcrossing. Table S3 enumerates the days of sporulation for each cycle as well as additional details regarding the culturing conditions for both versions of the 18-way population. After each round of sporulation, cultures were processed as detailed above with the following modifications: 5mL of SIS, 10mL of 1% Tween 20, and 5mL of SDS were used to kill vegetative cells. Tetrads were disrupted by bead milling at 1500 strokes/minute using the Geno Grinder 2000 for 25-45m. The contents of the tubes were mixed thoroughly with a pipette to ensure maximal recovery of cells from the bead slurry. The supernatant was then transferred to 50mL Falcon tubes, after which 500uL of YPDa was added back to the tubes, which were then vortexed at the highest setting briefly. The supernatant was transferred to the same 50mL Falcon tube. Cultures were harvested, washed, then resuspended in 5mL of YPDa. At this point, cells were carefully mixed by pipetting and then transferred to a 250mL Erlenmeyer flask with 7.5mL of YPDa. Spores were mated for 3-4h at 30°C at 40RPM, after which the presence of shmoos and/or zygotes was checked under the microscope. 12.5mL of YPDach was added to the mated cells, which were incubated O/N at 200RPM at 30°C. The next day, cells were transferred to 50mL Falcon tubes, 7mL of culture was mixed with an equal volume of 30% glycerol to make frozen stock, while the rest of the culture was spun down at 3000RPM for 5m. Cultures were washed twice, resuspended in 25mL of PA7, then transferred to a 250mL flask with 50uL of 100mg/mL ampicillin and sporulated at 30°C at 275RPM to initiate the next cycle of outcrossing. Following the twelfth cycle of sporulation followed by random mating, cells were transferred to 1L flasks with 187.5mL of YPDach and incubated O/N at 30°C at 200RPM. The following day, all 200mL of culture was mixed with an equal volume of 40% glycerol and frozen down at −80°C in a combination of 2mL cryotubes and 15mL Falcon tubes.

### Whole genome sequencing of the haploid founder strains

All 18 founder strains were sequenced using a combination of PacBio long read and Illumina short read technology. PacBio sequencing data was available from a previous study for 6 of the 18 strains (founder AB1-AB4, A7, and A9) (Yue *et al*. 2017), which was downloaded and reassembled using our pipeline so that all assemblies are directly comparable. The remaining strains were struck out onto YPD plates for 3d at 30°C, after which a single colony was inoculated into 50mL of YPDamp and incubated at 30°C at 200RPM O/N. DNA was extracted using the Qiagen G-tip DNA extraction kit. Purified genomic DNA was sheared using 24-gauge blunt needles. The resulting sheared gDNA samples were quality checked by a FIGE run at 134V O/N and concentrations were measured using Qubit. Sample were considered acceptable if the majority of gDNA was sheared to between 20kb and 100kb. In our hands carefully controlling the gDNA size distribution results in longer N50 PacBio reads which gives better *de novo* assemblies with less data. SMRTbell libraries were prepared and sequenced at the UCI Genomics High Throughput Facility using a PacBio RSII machine. The details of PacBio library creation for the purpose of *de novo* genome assembly are described in (Chakraborty *et al*. 2016). The average per-site coverage of the six previously sequenced strains was 365x as compared to 59x for the twelve strains sequenced in our hands, while the average PacBio read N50 for the previously sequenced strains was 5.73 kb as compared to 11.65kb for the strains sequenced by our lab.

Libraries for Illumina sequencing were made for all 18 founder strains. The same genomic DNA that had been used to prep the SMRTbell libraries was used to make Illumina libraries. Genomic DNA from the six remaining strains was prepared using the Qiagen G-tip kit as above. All genomic DNA was sheared to ~300-400bp using the Covaris S220 Focused Acoustic Shearer with the following settings: peak incident power (w) of 140, Duty Factor of 10%, Cycles per Burst of 200, Treatment time of 65s, temperature of 4°C, and water 12. Illumina compatible libraries were prepared using the NEBNext Ultra II DNA Library Prep kit along with the NEBNext Multiplex Oligos for Illumina (Index Primer Set 1) as per the manufacturer’s recommendations. Adaptor-ligated DNA was size-selected and PCR-enriched for five cycles, followed by clean-up of the PCR reaction using AMPure XP Beads as per the NEBNext Ultra II DNA Library Prep protocol. Sequencing was carried out using the Illumina HiSeq4000 with PE100 or PE150 reads (see Note S3). The average per-site coverage of the 18 founder strains was 290x with the lowest coverage being 186x (founder B11) and the highest coverage at 374x (founder AB4).

### Genome assembly

We assembled the PacBio reads using canu v1.7 (commit r8700; options: corMhapSensitivity=high, corOutCoverage=500, minReadLength=500, corMinCoverage=0, correctedErrorRate=0.105) (Koren *et al*. 2017). We generated hybrid assemblies using the PacBio and Illumina reads for the twelve strains for which we generated the PacBio reads. The PacBio reads from the six strains from (Yue *et al*. 2017) were too short to assemble with DBG2OLC, the hybrid assembler we use (Ye *et al*. 2016). The DBG2OLC hybrid assemblies were used to fill gaps in the corresponding canu assemblies using quickmerge, following the two steps merging approach (Chakraborty *et al*. 2016; Solares *et al*. 2018). The PacBio reads from Yue et al. were sequenced using an older chemistry of Pacific Biosciences (P4-C2) than our PacBio reads (P6-C4), so they required a different algorithm for optimal polishing than the assemblies created with the P6-C4 reads. Hence, we polished the P4-C2 based assemblies twice using Quiver and the P6-C4 based assemblies twice using Arrow (smrtanalysis v5.2.1). Finally, we polished all assemblies twice with the paired end Illumina reads using Pilon (Walker *et al*. 2014).

### BUSCO assessment

We estimated the number of fungi BUSCOs (n=290) in each polished assembly using BUSCO v3.0.2 (Waterhouse *et al*. 2018) (Table 2). For the augustus gene prediction step in BUSCO we used ‘saccharomyces_cerevisiae_S288C’ as the species option.

**Table 2.** Assembly statistics for the 18 sequenced founder strains.

| Strain | Assembly size (Mb) | Assembly N50 (kb) | Assembly QV | Assembly BUSCO (complete) | Total PacBio data (Mb) | PacBio read N50 (kb) | Total Illumina data (Mb) | Type of Illumina reads | x PacBio coverage | x Illumina coverage |
| --- | --- | --- | --- | --- | --- | --- | --- | --- | --- | --- |
| A5 | 12.13 | 757 | 48.8 | 0.990 | 737 | 11.10 | 3,396 | PE100 | 61.4 | 283.0 |
| A6 | 11.98 | 913 | 46.6 | 0.986 | 620 | 11.68 | 4,079 | PE150 | 51.7 | 340.0 |
| A7 | 12.62 | 901 | 66.2 | 0.990 | 3,979 | 5.97 | 2,775 | PE100 | 331.6 | 231.3 |
| A8 | 12.03 | 917 | 55.5 | 0.993 | 622 | 11.79 | 3,420 | PE100 | 51.9 | 285.0 |
| A9 | 12.53 | 571 | 53.0 | 0.993 | 5,845 | 5.10 | 2,769 | PE100 | 487.1 | 230.7 |
| A11 | 12.10 | 702 | 49.3 | 0.986 | 644 | 12.03 | 2,758 | PE100 | 53.7 | 229.8 |
| A12 | 12.00 | 795 | 45.5 | 0.993 | 613 | 11.83 | 3,563 | PE100 | 51.1 | 296.9 |
| B5 | 12.32 | 738 | 46.5 | 0.990 | 741 | 10.97 | 3,861 | PE100 | 61.7 | 321.8 |
| B6 | 12.10 | 772 | 44.9 | 0.990 | 401 | 11.67 | 2,906 | PE100 | 33.4 | 242.2 |
| B7 | 11.91 | 765 | 39.5 | 0.986 | 719 | 11.79 | 4,266 | PE100 | 59.9 | 355.5 |
| B8 | 11.99 | 802 | 66.0 | 0.990 | 741 | 12.00 | 3,054 | PE100 | 61.8 | 254.5 |
| B9 | 12.40 | 856 | 54.6 | 0.993 | 1,083 | 12.05 | 3,419 | PE100 | 90.2 | 284.9 |
| B11 | 12.12 | 789 | 55.0 | 0.990 | 765 | 12.13 | 2,236 | PE100 | 63.8 | 186.3 |
| B12 | 12.04 | 790 | 48.7 | 0.986 | 804 | 10.79 | 3,930 | PE100 | 67.0 | 327.5 |
| AB1 | 12.35 | 901 | 47.9 | 0.993 | 3,315 | 5.84 | 3,509 | PE150 | 276.3 | 292.4 |
| AB2 | 12.83 | 741 | 66.2 | 0.990 | 5,230 | 4.77 | 4,230 | PE150 | 435.8 | 352.5 |
| AB3 | 12.44 | 809 | 55.8 | 0.990 | 3,254 | 6.21 | 3,891 | PE150 | 271.2 | 324.2 |
| AB4 | 12.32 | 800 | 55.7 | 0.990 | 4,649 | 6.43 | 4,490 | PE150 | 387.4 | 374.2 |

\*Cells highlighted in blue represent founder strains that were previously sequenced using PacBio technology in (Yue *et al.* 2017)

### QV estimate

To estimate assembly error rate, paired end Illumina reads used in assembly polishing were mapped to the final assembly using bowtie2 (Langmead and Salzberg 2012). SNPs and small indels were identified using freebayes v0.9.21 (-C 10 -0 -O -q 20 -z 0.10 -E 0 - X -u -p 1 -F 0.75) (Garrison and Marth 2012a). To estimate the error rate, total bases due to SNPs and small indels (e) and the total number of assembly bases (b) with read coverage ≥3 were counted and qv was calculated as –10 X log(e/b) (Koren *et al*. 2017) (Table 2).

### Santa Cruz Browser Tracks

Assembled genomes were aligned to one another and the *SacCer3* reference genome using *ProgressiveCactus* (https://github.com/ComparativeGenomicsToolkit/cactus) (Paten *et al*. 2011a, 2011b). Santa Cruz Browser Track Hubs were created using the *hal2assemblyhub* script that is part of the *ProgressiveCactus* software (https://github.com/ComparativeGenomicsToolkit/Comparative-Annotation-Toolkit). The resulting SNAKE tracks are viewable at http://bit.ly/2ZrreUd. SNPs were identified between the founder strains using a generic GATK pipeline, with SNPs functionally annotated using SNPeff (Cingolani *et al*. 2012). Scripts to align the genomes and call SNPs in the founders are available here: https://github.com/tdlong/yeast_resource.

### Analysis of structural variants

We aligned each founder genome assembly to the s288c reference genome (GCA_000146055.2) using MUMmer v4.0 (Marçais *et al*. 2018) (nucmer --maxmatch -- prefix founder ref.fasta founder.fasta). To annotate the SVs, the delta alignment file for each strain was then processed with SVMU (commit e9c0ea1) (Chakraborty *et al*. 2019).

### Whole genome sequencing of the two base populations

The two base populations were deeply sequenced using Illumina technology. In total, 4mL of the 18F12v1 frozen stock was thawed at RT, pelleted at 3000 RPM for 5m, and resuspended in 20mL of YPDamp. This was followed by incubation at 30°C for 3.5h at 275RPM. Genomic DNA was extracted using the Qiagen DNeasy kit. The genomic DNA was sheared using the Covaris S220 as above and Illumina compatible libraries were prepared using the NEBNext Ultra II DNA Library Prep kit as above. The NEBNext and libraries were pooled and sequenced on the HiSeq4000 using PE100 reads. The NEBNext libraries were sequenced at a mean per-site coverage of 3270x.

For 18F12v2, similarly to 18F12v1, 4mL of frozen stock was thawed at RT, pelleted at 3000RPM for 5m, and resuspended in 20mL of YPDamp, followed by incubation at 30°C for 3.5h at 275RPM. Genomic DNA was extracted using the Qiagen G-tip kit. Nextera libraries were prepped for 18F12v2 by following the standard Nextera protocol with slight modifications. Tagmentation reactions were carried out in 2.5uL reactions for 10m at 55°C. Reactions were stopped by adding SDS to a final concentration of 0.02% followed by incubation at 55°C for 7m. Samples were immediately transferred to ice. Limited cycle PCR was carried out to add two unique barcodes to each library to enable dual index sequencing. This avoids the problem of barcode switching when *N* i7 and *M* i5 barcodes are used to create *MN* combinations. The KAPA HiFi Ready Mix (2X) was used in conjunction with the KAPA forward and reverse primers to amplify tagmented libraries in 25uL total volume. Thermocycling parameters consisted of 3m at 72°C, 5m at 98°C, followed by 15 cycles of 10s at 98°C, 30s at 63°C, and 30s at 72°C, with a hold of 72°C for 5m at the end. PCR reactions were cleaned up using AMPure XP Beads (Beckman Coulter, Inc.) and libraries quantified by Qubit. Sequencing was performed on the HiSeq4000 using PE150 reads. 18F12v2 libraries received 2226x coverage.

### Whole genome sequencing of the first two meiotic generations of the second base population

For 18F1v2 and 18F2v2, ~1mL of frozen stock was thawed at RT, spun down at 7500 RPM for 5m in microcentrifuge tubes, resuspended in 1mL of YPDamp, transferred to a 250mL flask with 19mL of YPDamp and incubated at 30°C for 3.5h at 275RPM. Genomic DNA from both samples was extracted using the Qiagen G-tip kit. Nextera libraries were prepped by following the Nextera flex protocol using 1/5^th^ reactions with slight modifications. Limited cycle PCR was carried out using the KAPA HiFi Ready Mix (2X) as detailed above to add barcoded Illumina-compatible adapters in 12.5uL reactions. Thermocycling parameters consisted of 3m at 72°C, 3m at 98°C, followed by 12 cycles of 45s at 98°C, 30s at 62°C, and 2m at 72°C, with a hold of 72°C for 1m at the end. Proteinase K was added to each reaction (50ug/mL final concentration) to digest the polymerase. Samples were incubated for 30m at 37°C and 10m at 68°C. Reactions were cleaned up using the SPB beads provided with the Nextera flex kit. Sequencing was performed as above using PE100 reads. 18F1v2 received 98x coverage while 18F2v2 received 73x coverage.

### Whole genome resequencing of recombinant haploid clones

Ten haploid recombinant clones (5 of each mating type) were isolated from each of the two base populations. 18F12v1-derived haploids were generated by sporulating an O/N culture of the 18F12v1 population in 2mL of PA7 in a 10mL culture tube at 30°C for 3d. Spore isolation and dispersal were carried out as detailed above for the creation of 18F12v2 with 15m of bead milling to disperse spores. Spores were plated at low density onto YPD plates and incubated for 2d at 30°C. One of the YPD plates was then replica plated onto four different plates: YPD with hyg, YPD with cloNAT, YPD with mate-type tester 1, and YPD with mate-type tester 2. Five haploids of each mating type were inoculated into YPD O/N. Genomic DNA was extracted using the Qiagen DNeasy kit and Nextera libraries prepared as above. Libraries were sequenced on a HiSeq4000 using PE100 reads to a mean per-site coverage of ~32x.

18F12v2-derived haploids were generated by sporulating an O/N culture of the 18F12v2 population in 4mL of PA7 in a 24 deep-well plate at 30°C at 275RPM for 3d. Spore isolation and dispersal were carried out as detailed above for the creation of 18F12v2 with 20m of bead milling to disperse spores. Spores were plated at low density onto YPD plates and incubated at 30°C for 3d. 96 single colonies were then transferred into a 96 deep-well plate with YPDamp using sterile toothpicks. After O/N incubation at 30°C, 200uL of culture from each well was transferred to a 96 shallow plate and pinned YPD plates with either cloNAT, hyg, mate-type tester 1, or mate-type tester 2 using a 48-well replicator tool. The source plate was covered with an adhesive membrane and stored at 4°C. The mate typing plates were incubated at 30°C for 2d, after which five haploids of each mating type were transferred from the original source plate to 1.5mL eppendorfs and genomic DNA was extracted using the Qiagen DNeasy kit. Nextera libraries were prepared as above. Libraries were sequenced on a HiSeq4000 using PE150 reads to a mean per-site coverage of ~60x.

### Haplotype calling in Illumina resequenced MPPs and recombinant haploid clones

De-multiplexed fastq files were used in analyses. Detailed scripts/software versions to reproduce our analysis are located at https://github.com/tdlong/yeast_resource.git. Briefly, reads were aligned to the *sacCer* reference genome using *bwa-mem* and default parameters (Li and Durbin 2009; Li 2013). We maintain two SNP lists, a set of known SNPs in the strains obtained from a GATK pipeline that only considers the isogenic founders, and a subset of those SNPs that are well-behaved (i.e., frequency of the REF allele close to zero or one in all founder lines, pass GATK qualify filters, etc.). The list of well-behaved SNPs that are polymorphic in the founders can be used to speed-up subsequent steps, where we sometimes examine hundreds of samples, since only variants polymorphic among the founders need be considered when working with samples from a synthetic population (except when calling newly arising mutations). *samtools mpileup* (Li *et al*. 2009; Li 2011) and *bcftools* (Narasimhan *et al*. 2016) are used to query well-behaved known SNPs. We have no interest in calling genotypes, but instead simply output the frequency of the REF allele in each sample at each location (output=SNPtable). In a separate analysis *freebayes* (Garrison and Marth 2012b), *vcfallelicprimitives* (https://github.com/vcflib/vcflib), and *vt normalize* (Tan *et al*. 2015) are used to call all SNPs, and the SNPs not in our list of known SNPs considered candidate new mutations.

We have developed custom software to infer the frequency of each founder haplotype at each location in the genome in pooled samples using the SNPtable as input and the *haplotyper.limSolve.code.R* script in the github archive. This same algorithm can also be used without modification to infer genotypes in recombinant haploid clones. Briefly, we slide through the genome in 1kb steps considering a 60kb window for each step. For all SNPs in the window we calculate a Gaussian weight such that the 50 SNPs closest to the window center account for 50% of the sum of the weights. We then consider F founders and use the *lsei* function of the *limSolve* package (*limSolve: Solving Linear Inverse Models*, R package 1.5.1) (Meersche *et al*. 2009) in *R* to identify a set of F mixing proportions (each greater than zero and summing to one) that minimize the sum of the weighted squared differences between founder haplotypes and the observed frequency of each SNP in a pooled sample. That is, for a N SNP window we call *lsei* with the following parameters: A=N*F matrix of founder genotypes, B=N*1 vector of SNP frequencies in a pooled sample, E=F*1 vector of 1’s, F=1, G=F*F identity matrix, H= F*1 vector of 0’s, and Wa=N*1 vector of weights. Finally, for windows where the i^th^ and j^th^ founders have near indistinguishable haplotypes, implying the sum of the two mixing proportions are correct, but not individual estimates, we estimate the haplotype frequency as half the sum of the two mixing proportions. This method of accounting for indistinguishable haplotypes is regional and is generalized to more than two near identical founders and multiple such sets.

### Validation of the haplotype caller

In order to validate our haplotype-calling algorithm, we identified 70,478 SNPs private to a single founder strain (excluding those present in founders merged due to high sequence similarity). The haplotype caller was run on 18F12v2 using the full coverage data (i.e., 2230X) or down-sampled 18F12v2 to simulate a more typical poolseq re-sequencing depth (typical applications using the MPPs are likely to sequence hundreds of experimental units to ~20-60X). For each private SNP, the frequency of the SNP in the full coverage data was estimated and the founder harbouring that SNP identified. Since the sequence depth of the non-downsampled population is 2230X, the frequency of each private SNP is measured very accurately. We then infer the frequency of the founder haplotype harbouring the private SNP at the position closest to the private SNP, in both the full coverage and each downsampled population. The error rate associated with the haplotype caller is the absolute difference between frequency of each private SNP and the founder haplotype harbouring it.

In our examination of the relationship between haplotype and SNP frequency estimates (Figure 4) we identified and removed 91 outlier SNPs among the 71,301 private SNPs. These SNPs were identified as private SNPs whose frequency was more than 5% different than the frequency of the haplotype harbouring it in the full dataset, while exhibiting flanking private SNPs in the same founder whose frequency agree with the founder frequency. We believe these outlier SNPs are cases where that particular SNP in a pooled sample cannot be aligned to the reference genome very accurately. It is noteworthy that it is more difficult to identify poorly performing SNPs that are not private to a single founder, and such SNPs likely hurt haplotype inference methods.

**Figure 4.**
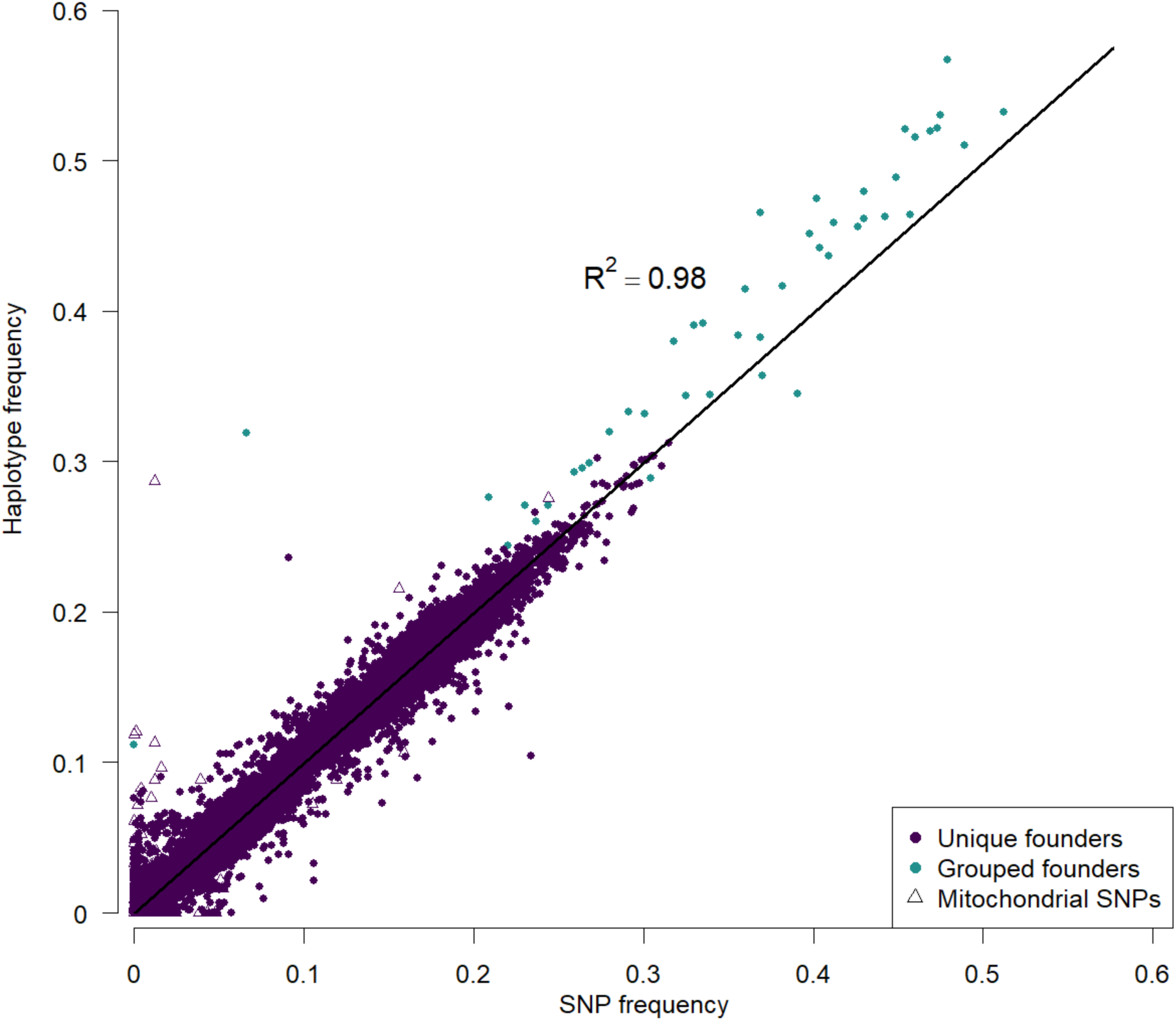
The frequency of SNPs private to a single founder are highly correlated with the estimated haplotype frequencies at these SNPs in 18F12v2. As the frequency of a private SNP should be equal to the corresponding haplotype frequency, this measure provides a benchmark with which the accuracy of our haplotype caller can be measured. Cyan points represent founders that were pooled when estimating haplotype frequencies (“grouped founder”) due to the high degree of sequence similarity between their genomes. Triangles represent mitochondrial SNPs, which, together with SNPs private to pooled founders, represent the bulk of the major outliers. The coefficient of determination was calculated by regressing haplotype frequency onto SNP frequency, excluding SNPs from grouped founders and mitochondrial SNPs.

### Delineating haplotype blocks in recombinant haploid clones

The haplotype caller is primarily used to estimate the frequency of founder haplotypes at different positions in the genome in a DNA pool from a segregating population, but it can also be run on DNA obtained from a haploid or diploid clone. In a haploid clone the haplotype caller should return a haplotype frequency of close to 100% for one of the founder haplotypes for much of the genome, with sharp transitions between founder states near recombination breakpoints. In depicting the haplotypic structure of haploid clones we classify genomic regions at which the inferred haplotype frequency of a single founder (or multiple indistinguishable founders) is less than 95% as having an ‘unknown’ haplotype (these unknown intervals typically being associated with state transitions). We also observe intervals in which several founders are indistinguishable from one another (due to insufficient SNP divergence between the founders in these window). We could sometimes resolve these intervals to a single founding haplotype when flanking haplotypes were unambiguously called as derived from the same single founder. Custom R scripts were used for these analyses as well as to calculate the length of haplotype blocks in haploid clones. Haplotype block sizes were inferred by finding the positional difference between the beginning and end of runs of the same haplotype.

### Data and reagent availability

Strains and plasmids are available upon request. All genome sequencing data and assemblies have been deposited into public repositories. Sequence data generated for the two base populations (18F12v1, 18F12v2, 18F1v2, and 18F2v2) as well as the recombinant haploid clones are available in the Short Reads Archive under the bioproject PRJNA551443 in accessions SRX6465384 to SRX6465405 and SRX6983898 to SRX6983899. All PacBio and Illumina data generated for the 18 founding strains is also available in the Short Reads Archive under the bioproject PRJNA552112 in accessions SRX6380915 to SRX6380944. Detailed scripts/software versions to reproduce our analysis are located at https://github.com/tdlong/yeast_resource.git.

## Results and Discussion

### Recovery of hyg^r^ and insertion of dominant selectable markers for high-throughput diploid selection

We further engineering a subset of the yeast SGRP resource strains (Cubillos *et al*. 2009) to serve as founders for an 18-way synthetic population. We first recovered the *hyg^r^* marker used to delete the *HO* gene in the haploid SGRP strains. Previous work (McDonald *et al*. 2016) replaced *YGR043C*, a pseudogene that is physically close to the mating type locus, with dominant selectable markers in order to facilitate high-throughput selection of diploids after mating. We echoed that approach here by replacing YGR043C with *NatMX4* in 15 *Mat a* (or “A”) founders and with *HphMX4* in 15 *Mat α* (or “B” founders). The presence of these cassettes confers resistance to the antibiotics nourseothricin and hygromycin B, respectively, enabling the selection of doubly resistant diploids. All newly engineered strains are given in Table 1.

### de novo assembly of high-quality reference genomes for the 18 founding strains

We generated *de novo* genome assemblies for the founders used to create our MPPs using a hybrid sequencing strategy detailed in (Chakraborty *et al*. 2016) that involves using a combination of long-read (PacBio) and short-read (Illumina paired-end) sequencing technology. The *de novo* assemblies allow us to reliably identify structural variants while the overall assembly has a low per base pair error rate. We assembled 58.9X PacBio reads on average (33-90X) for 12 of the founder strains, and re-assembled the other six strains using publicly available shorter length 364.9X PacBio reads on average. Despite the different number and chemistry of PacBio reads used in assembling the genomes, all of our assemblies are highly accurate (average qv = 52.5) and show comparable contiguity. For example, the average contig N50 of our assemblies is ~800Kb (N50 = 50% of the assembly is contained within sequences of this length or longer), indicating that the majority of the chromosomes are represented as single contigs (Table 2). Examination of 290 conserved fungal single copy orthologs (Benchmarking Single Copy Orthologs or BUSCO) show that completeness (~99%) of all our assembled genomes is comparable to the reference S288C assembly (99%).

We aligned the assemblies to one another and represent them as Santa Cruz genome browser tracks (http://bit.ly/2ZrreUd). These tracks have utility when looking for candidate causative variants in small regions of genetic interest. The large amount of genetic diversity sampled by the founders can be illustrated by zooming in on regions such as that shown in Figure 2, which highlights the numerous alleles segregating at a gene implicated in many genetic mapping studies in budding yeast, the highly pleiotropic *MKT1*. *MKT1* influences several cellular processes including the DNA damage response, mitochondrial genome stability, drug resistance, and post-transcriptional regulation of *HO* (Dimitrov *et al*. 2009; Ehrenreich *et al*. 2010; Tkach *et al*. 2012; Kowalec *et al*. 2015a). Studies have found that different alleles of *MKT1* can differentially affect several phenotypes, including mitochondrial genome stability and drug resistance. Variation at this gene amongst our founders includes ten nonsynonymous SNPs and thirty-four synonymous SNPs. Of the ten nonsynonymous SNPs, six are predicted to change the secondary structure of the protein. Taking into account only nonsynonymous SNPs, there are seven different alleles segregating amongst the founders (all segregating in our 18F12v2 MPP).

**Figure 2.**
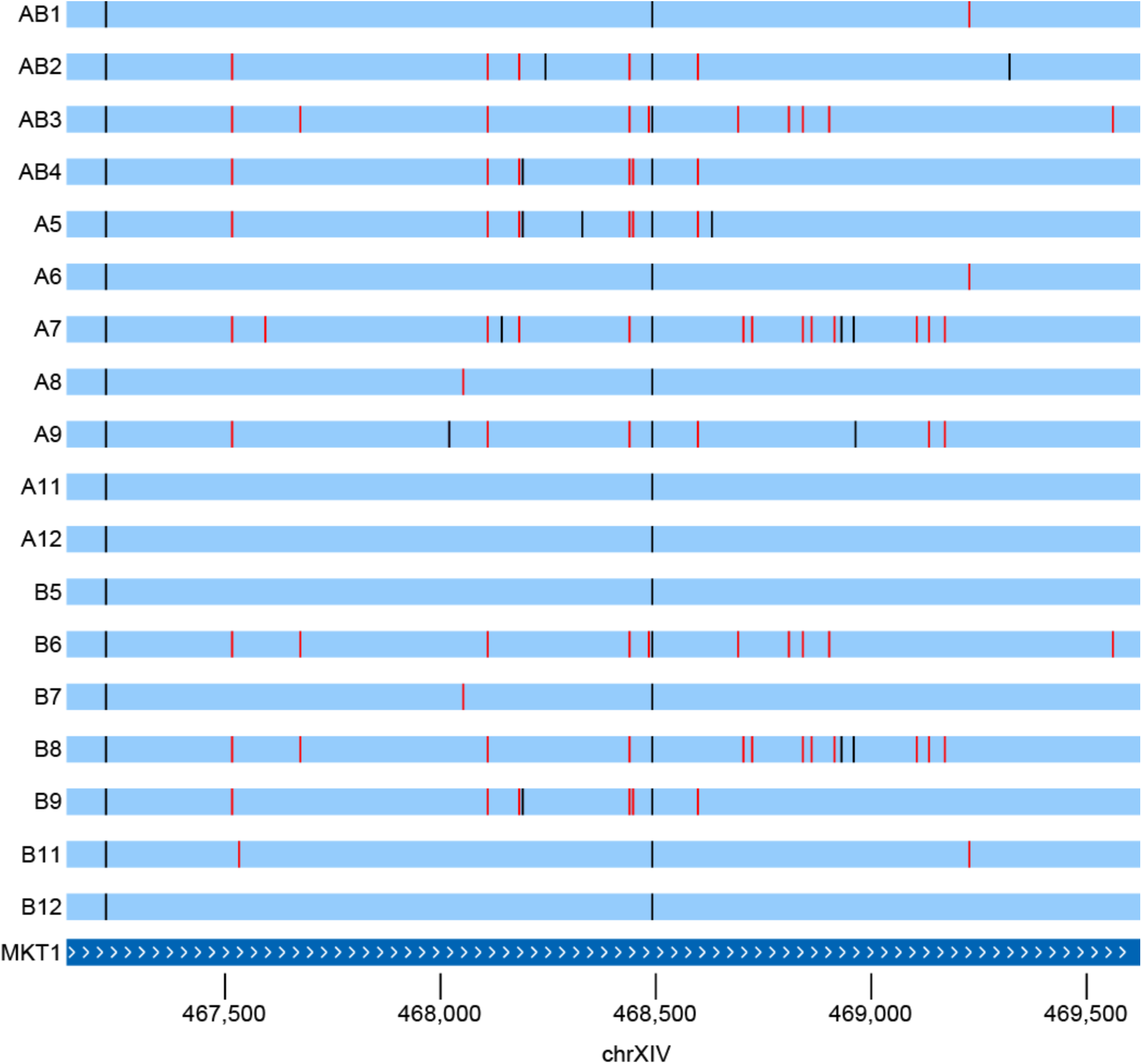
Many alleles of the highly pleiotropic *MKT1* gene are segregating amongst the founder strains, highlighting the potential of uncovering complex allelic series using populations derived from these strains. Seven of these alleles are differentiated by nonsynonymous SNPs, of which six are predicted to be segregating in 18F12v2. Vertical red lines are synonymous SNP differences from the reference S288C strain, and black bars nonsynonymous SNPs.

The genome browser tracks are also useful for visualizing structural variants such as those shown in Figure 3, which highlights a large (>1kb) deletion in the vacuolar ATPase *VMA1* (Figure 3A) present in half of the founders. Previous work has shown that the deleted region encodes a self-splicing intein, PI-SceI, a site-specific homing endonuclease that catalyzes its’ own integration into inteinless alleles of *VMA1* during meiosis (Gimble and Thorner 1992). This selfish genetic element has been shown to persist in populations solely through horizontal gene transfer and is present in many species of yeast. Perturbation of *VMA1* itself has been shown to influence both replicative and chronological lifespan, resistance to metals, as well as oxidative stress tolerance (Kane 2007; Ruckenstuhl *et al*. 2014).

**Figure 3.**
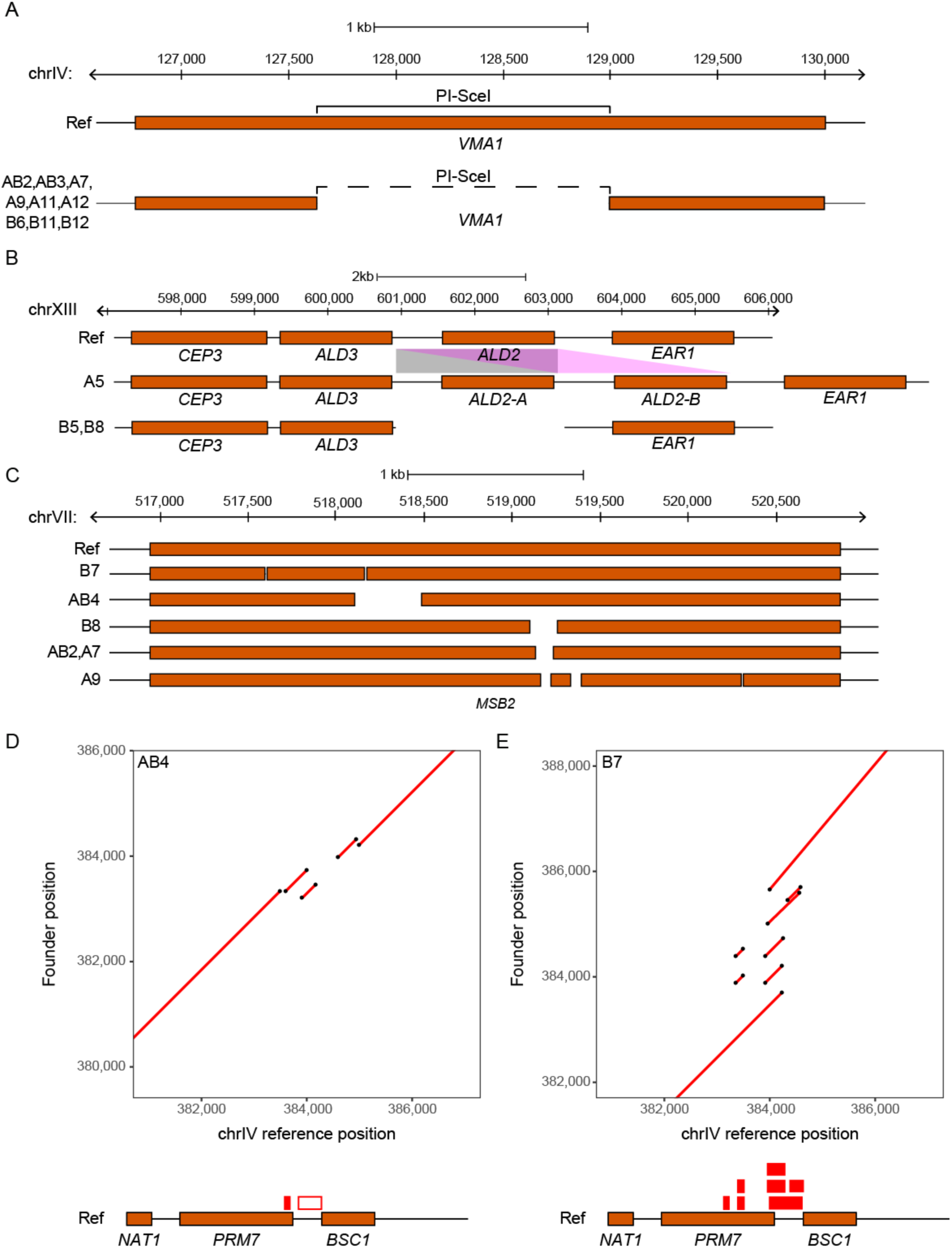
Combining contiguous long-read sequencing with accurate short-read data enables the detection of structural variants such as those depicted above. In (**A**), a large (>1kb) deletion within a vacuolar ATPase (*VMA1*) is present in half of the strains used in this study. This deletion directly overlaps the self-splicing intein, PI-SceI. Copy number variants of *ALD2*, an aldehyde dehydrogenase, were detected (**B**) and include a duplication of this gene in founder A5 (represented as *ALD2-A* and *ALD2-B*) as well as its’ deletion in founders B5 and B8. In (**C**), multiple deletions of different lengths in the osmosensor *MSB2* were detected in multiple founder strains. Dotplots of a structurally complex region on Chromosome IV are shown for founders AB4 (**D**) and B7 (**E**). These plots show alignments of regions from the founder strains (depicted on the y axis) with the corresponding region from the S288C reference strain (depicted on the x axis). The red boxes present above the genes in the reference strain map duplications (solid boxes) and deletions (empty boxes) detected in each founder strain to the corresponding reference sequence. In all panels, the reference strain S288C (aka ‘Ref’) is used to highlight the various arrangements of SVs present in the founder strains.

In addition to large deletions, copy number variants (CNVs) can also be found, such as that shown in Figure 3B, in which the cytoplasmic aldehyde dehydrogenase, *ALD2*, is duplicated in founder A5. Conversely, this gene has been deleted in founders B5 and B8. *ALD2* has been shown to be involved in the osmotic stress response as well as the response to glucose exhaustion (Navarro-Aviño *et al*. 1999). A more structurally complex region was identified on Chromosome VII (Figure 3C) at which multiple different deletions (ranging from ~50bp to >300bp) were found to occur at *MSB2*, an osmosensor involved in the establishment of cell polarity (O’Rourke and Herskowitz 2002; Cullen *et al*. 2004). Null alleles of *MSB2* have been shown to have decreased chemical resistance.

One of the most structurally complex regions we identify contains ~2kb of repetitive sequence and is present on chromosome IV (Figure 3D and E as well FigureS1) at which multiple different deletions (ranging from ~80bp to >500bp) as well as duplications occur in multiple founders within the *PRM7* and *BSC1* genes. Due to the highly complex nature of the variation present, this region is represented as a series of dot plots, with two founders highlighted in the main text (Figure 3D and E). Dot plots of this region in all founder strains are shown in FigureS1. A previous study demonstrated that although two distinct genes (*PRM7* and *BSC1*) are present in S288C, a combination of small deletions and point mutations in another yeast strain (W303) have caused the STOP codon to be absent from *BSC1*, leading to the read-through transcription of a new gene that encompasses sequence from both *PRM7* and *BSC1* as well as the intergenic region between them (Kowalec *et al*. 2015b). This gene, *IMI1*, was shown to affect mtDNA stability as well as intracellular levels of reduced glutathione (GSH).

The above regions highlight the utility of our *de novo* genome assembly approach, as deletions and CNVs of this scale would be difficult to detect via the usual method of aligning short reads to a reference genome. But if an investigator mapped a QTL to one of these genes they would certainly want to know about the existence of the segregating structural variation.

Despite the large amount of natural variation present amongst the founders in general, some of the founders were found to be genetically very similar to one another (AB3/B6) Figure S2 and shown in Table S4), having less than 200 pairwise SNP differences. This lack of divergence makes this set of founders difficult to distinguish from one another for much of the genome and as a result we collapse them for subsequent analyses (despite that fact that a subset of these 200 differences could be functional). Three additional founders (A11/A12/B11) were also found to be highly genetically similar to one another, with, on average, less than 2,000 pairwise SNP differences. These differences are concentrated in a small number of regions, making these three founders distinguishable for these regions (but indistinguishable for much of the remainder of the genome). We keep these strains separate for downstream analyses.

### Creation of two 18-way highly outcrossed populations

MPPs created using multiple rounds of recombination can significantly increase the resolution of genetic mapping studies by virtue of haplotypes sampled from these population having a greater number of genetic breakpoints. Furthermore, multiple founders results in high levels of standing variation present in the MPP. These two features result in populations that more realistically mimic natural outbred diploid populations, and samples more functional alleles and haplotypes from the species as a whole than a two-way cross. With these goals in mind, we constructed a large, genetically heterogenous population by crossing 18 different founder strains (each strain being derived from the SGRP (Cubillos *et al*. 2009)). The 18 founder strains were chosen to represent a broad swathe of the natural diversity of the species and belong to diverse phylogenies, including: Wine/European, West African, North American, Sake, and Malaysian (see Table 1). It is also noteworthy that founder strains A1-4 and B1-4 are the same four strains using in (Cubillos *et al*. 2013), and were introduced into the population as both *Mat a* and *Mat α* mating types. We created two versions of our 18-way MPP. In both cases a full diallele cross was used to create all 121 unique diploid genotypes from 11 *Mat a* and 11 *Mat α* strains (see Figure 1A and B). All 121 diploid genotypes were combined and the resulting population was taken through 12 rounds of sporulation followed by random mating to break up linkage disequilibrium. Previous work has shown that 12 rounds of random recombination breaks up haplotype blocks to the point where additional outcrossing does not significantly decrease LD (Parts *et al*. 2011). For brevity, the two different outcrossed populations will be referred to as 18F12v1 and 18F12v2, respectively, throughout the rest of this manuscript. The version 1 MPP differed primarily from version 2 in that the 121 diploid genotypes obtained from the diallele were directly combined and sporulated *en masse* (version 1; Figure 1A) to create the MPP, as opposed to being individually carried through sporulation and spore disruption before being combined (version 2; Figure 1B). Furthermore, due to a technical artefact during the 12 rounds of outcrossing, 18F12v1 is cross-contaminated with the 4-way F12 population from (Cubillos *et al*. 2013) which contains a functional URA3 gene. As a result, 18F12v1 MPP is of limited utility for experiments that require uracil auxotrophy, and 18F12v2 is the current primary focus of work in our lab.

### Development of an algorithm for accurately inferring haplotype frequencies

In QTL mapping experiments using MPPs it is often advantageous to map QTLs back to founder haplotypes. In experiments derived from a two way cross between isogenic founders genotyping SNPs accomplishes this, but with multiple founders parental haplotypes have to be inferred in recombinant offspring (Mott *et al*. 2000). In a similar manner when MPPs are used as a base population and genetic changes detected following an experimental treatment it is often of value to examine changes in haplotype frequency (as done in (Burke *et al*. 2014) and reviewed in (Barghi and Schlötterer 2019)). We developed a sliding window haplotype caller that can be used in the situation when the founder haplotypes are known and apply it to both single haploid clones and pools consisting of millions of diploid individuals. This haplotype caller differs from other widely used callers (Long *et al*. 2011; Kessner *et al*. 2013) in that it acknowledges that in some windows pairs of founders are poorly resolved or indistinguishable and relies solely on read counts at known SNP positions in both founders and recombinant populations.

To benchmark the haplotype-calling algorithm, we compared the frequency of SNPs private to a single founder to the haplotype frequency of the same founder for the interval closest to the SNP location in the 18F12v2 base population. Since this base population is sequenced to 2226X we initially wished to look at the error in the haplotype estimate at full coverage where the sampling variation on the SNP frequency estimate is quite low (proportional to 1/sqrt(2226) or <2%). For the high coverage base population regions showing large difference between SNP and haplotype frequency estimates likely represent instances where the haplotype caller breaks down, since we attempted to remove SNPs whose frequencies are poorly estimated. Figure S3, depicting the absolute difference in SNP versus haplotype frequency differences, shows that haplotype and SNP frequencies generally agree with one another with average and median error rates of 0.4% and 0.2%, respectively (below the sampling error of SNP frequency).

Of course, typical experiments employing these base populations will sample the population following some treatment, and comparing haplotype frequencies in control versus treated samples. Although the 18F12v2 base population is sequenced to 2226X, it would be cost-effective if we could infer haplotype frequencies from pooled samples sequenced to much lower coverage. To determine the accuracy of our haplotype estimates as a function of sequencing coverage the 18F12v2 was down-sampled 50-fold and 100-fold, which corresponds to poolseq datasets of ~40X and ~20X respectively. We then estimated relative haplotype frequency error rates as a function of sequence coverage (Figure S3b) and absolute error rates as a function of coverage and genomic location (Figure S4). It is apparent that the error rate is an increasing function of decreasing coverage, but for much of the genome the absolute error in haplotype frequency estimate is actually lower than the binomial sampling errors associated with directly estimating SNP frequencies at the same coverage (*i.e.*, at 20-40X coverage binomial sampling errors on frequency are >10%). It is also apparent that the average error rate is likely driven by a few regions where the haplotype caller struggles; these are presumably regions with poor divergence between founders in the window examined. Overall the mean (median) error rates on haplotype frequency estimates are low, 1.3% (0.8%) at 20X and 1% (0.6%) at 40X, respectively.

### Characterization of 18F12v1 and 18F12v2 base populations

18F12v1 and 18F12v2 were subjected to high coverage whole-genome sequencing to both characterize their population structure and to establish a baseline for future mapping studies. Figures 5 and 6 show the inferred sliding window haplotype frequencies for 18F12v1 and 18F12v2, respectively, while Table 3 shows the mean per founder haplotype frequencies genome-wide. One trend that is evident is that in both the 18F12v1 and 18F12v2 MPPs a small number of founders are over-represented. In order to identify the origin of this bias, at least for 18F12v2, the first two meiotic generations of 18F12v2 were sequenced (Figures S5 and S6; Table S5). Despite having an initially more balanced population after the first round of random mating, a few strains quickly became disproportionately over-represented. One possible explanation for this is that a few founding haplotypes were selected for early in the twelve rounds of intercrossing. Figure S7 provides suggestive evidence that this may have been the case, as the frequency of hapotypes derived from founder A5 increase genome-wide after the second round of meiosis. The latter half of chromosome XIII (from founder A5) emphasizes this point as it was very highly selected for initially. Another potential source of bias was the pooling strategy, which was done using optical density as a proxy for cell numbers. This may have resulted in an uneven distribution of founders in the initial pool. Nonetheless, after 12 rounds of random mating, deep sequencing of 18F12v2 revealed that haplotypes from all founding strains were present in the population at a detectable frequency (Table 3). Specifically, haplotypes from ten or more founders were detected as segregating in over 99% of the genome in 18F12v2 and close to 98% of the genome in 18F12v1. Furthermore, 18F12v2 was verified to be auxotrophic for uracil, facilitating future manipulations for downstream analyses.

**Figure 5.**
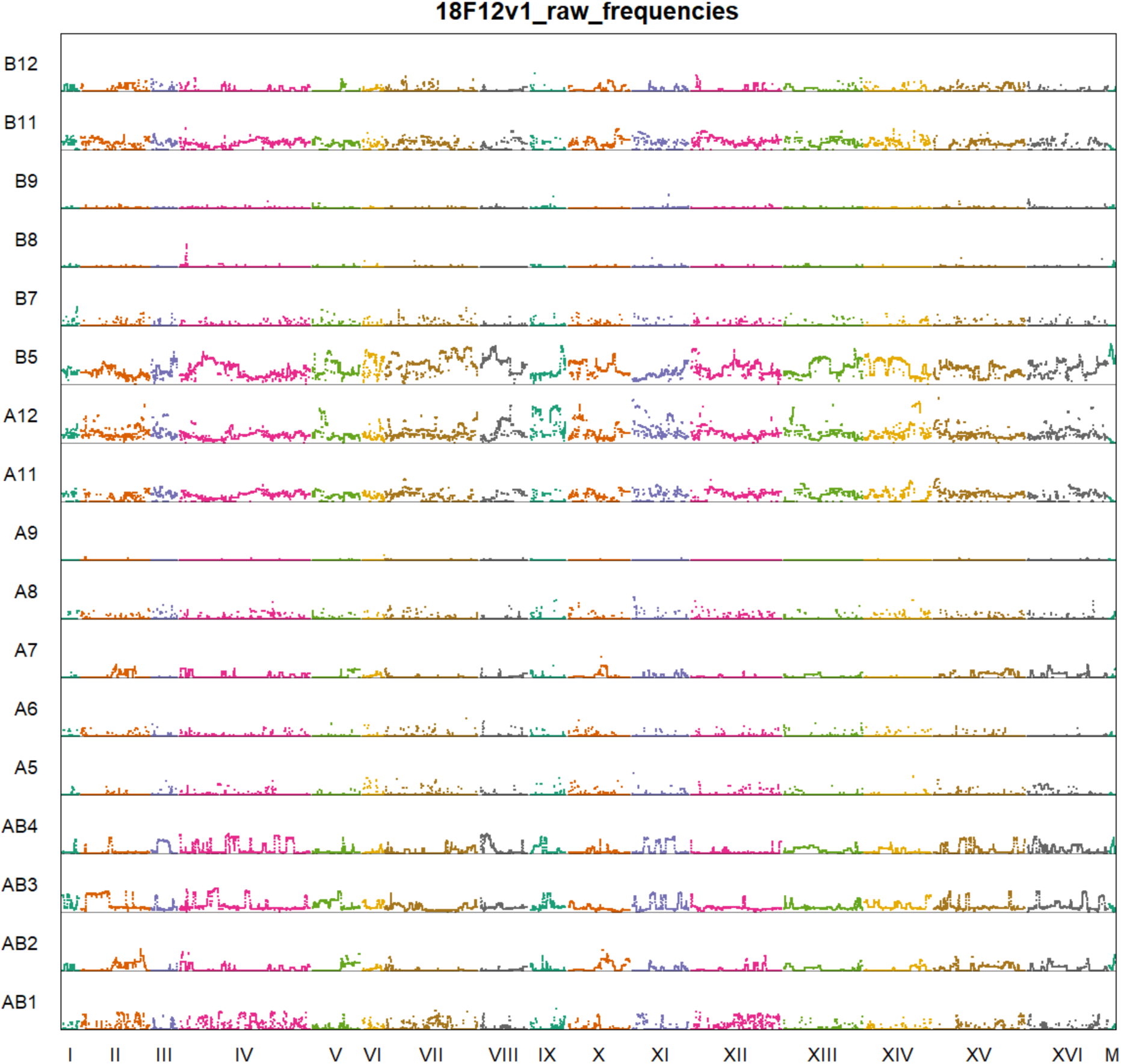
Genome-wide haplotype frequencies for 18F12v1.

**Figure 6.**
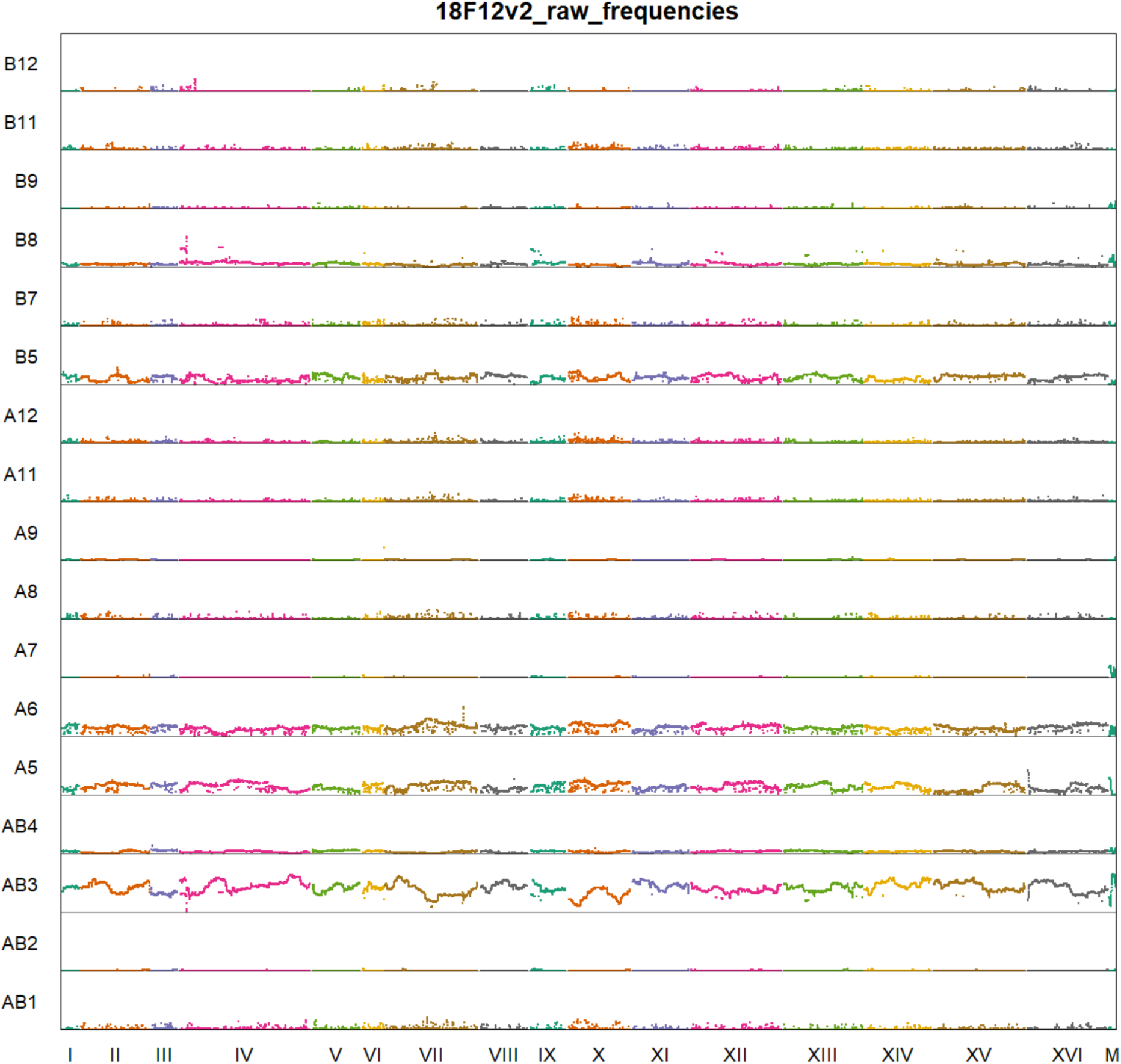
Genome-wide haplotype frequencies for 18F12v2.

**Table 3.** Mean haplotype frequencies in 18F12v1 and 18F12v2

| Founder | 18F12v1_frequency | 18F12v2_frequency |
| --- | --- | --- |
| AB1 | 4.4% | 1.2% |
| AB2 | 3.8% | 0.5% |
| AB3 | 10.6% | 41.4% |
| AB4 | 5.8% | 3.3% |
| A5 | 1.2% | 14.0% |
| A6 | 0.8% | 14.3% |
| A7 | 2.4% | 0.4% |
| A8 | 1.2% | 0.7% |
| A9 | 0.1% | 0.5% |
| A11 | 9.9% | 1.2% |
| A12 | 18.1% | 1.7% |
| B5 | 27.1% | 11.3% |
| B7 | 1.1% | 1.0% |
| B8 | 0.5% | 5.7% |
| B9 | 0.9% | 0.8% |
| B11 | 9.8% | 1.3% |
| B12 | 2.4% | 0.6% |

### Characterizing the recombination landscape of 18F12v1- and 18F12v2-derived segregants

In order to further characterize 18F12v1 and 18F12v2, ten haploid segregants were generated from each diploid population and subjected to whole-genome sequencing. The complex structure of these populations is highlighted in Figure 7. The mean (median) size of haplotype blocks in 18F12v1-generated segregants was 103kb (66kb) while the mean (median) size of haplotype blocks in 18F12v2-generated segregants was 106kb (66kb) (Figure S8). The mean number of discrete haplotype blocks in 18F12v1-generated segregants was 106 as compared to 104 in 18F12v2-generated segregants. A previous study (Cubillos *et al*. 2013) found that twelve rounds of meiosis in a yeast 4-way cross resulted in a median block size of 23kb with 374 discrete haplotype blocks. Some of the failure to obtain the smaller block sizes and more numerous discrete blocks of this previous study may be due to undetectable recombination events occurring within haplotypes over-represented in our populations. Another possibility is that, due to the large number of founding haplotypes, recombination events were missed in regions at which multiple founding strains are highly genetically similar. It is also possible that some of the founding strains used in this study have relatively low natural recombination rates.

**Figure 7.**
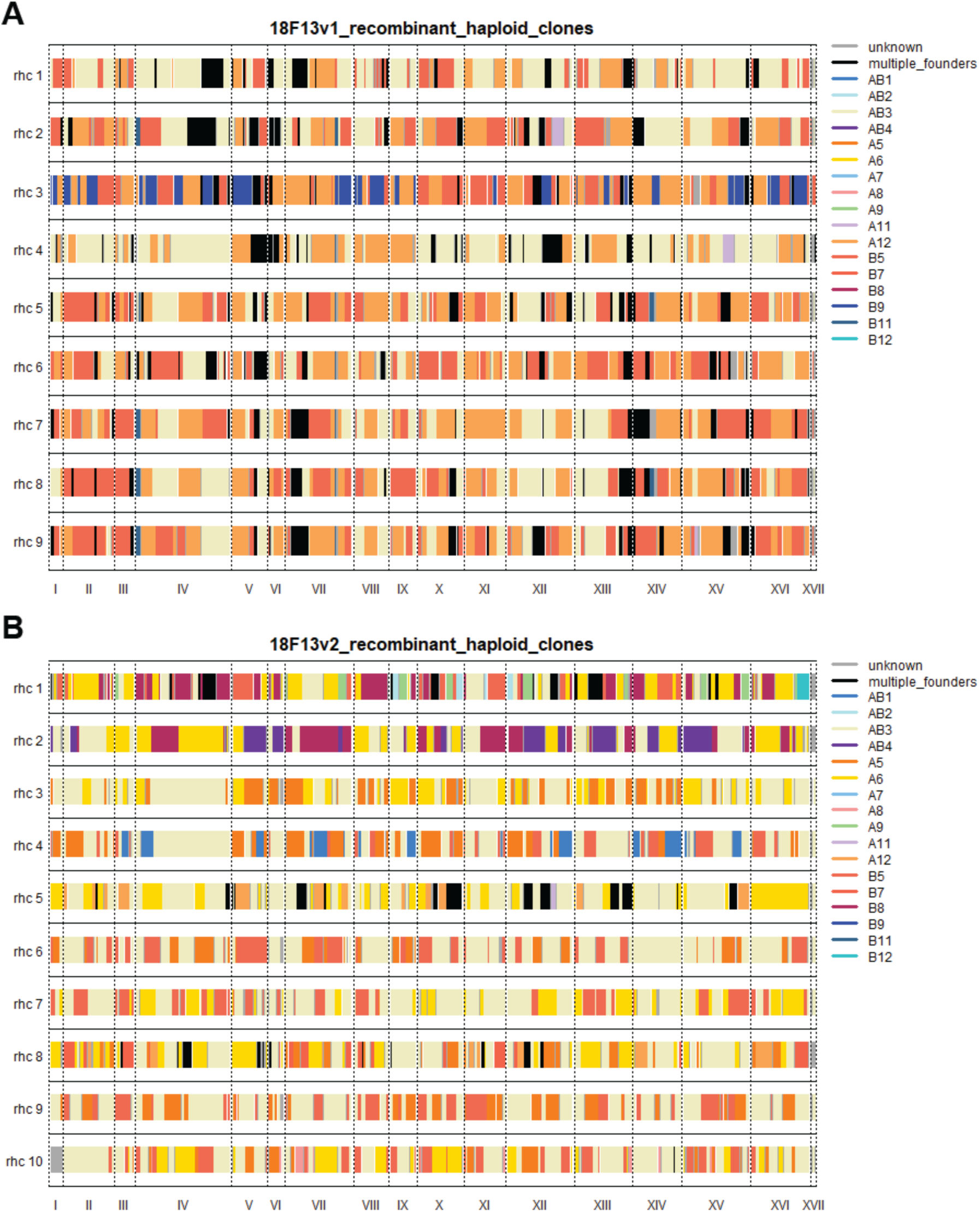
Haploids derived from 18F12v1 (**A**) and 18F12v2 (**B**) were isolated and sequenced, providing a glimpse into the recombinogenic landscape and haplotype diversity present within these populations.

To highlight the diversity present in the two outbred populations, a close-up view of inferred haplotypes in segregants derived from each population at chromosome X is shown in Figure S9. Regions in which the founding haplotype is unknown tended to occur at the transitions between haplotypes (see Note S4) and are a mean (median) length of 7.8kb (6kb) in 18F12v1-derived segregants and 7.5kb (6kb) in 18F12v2-derived segregants. Also noticeable is, at least for this chromosome, the larger amount of variation segregating in 18F12v2 (Figure S9B).

## Conclusion

The paradigm of utilizing pairwise crosses to dissect the genetic basis of complex traits has enjoyed much success in diverse model organisms. However, such studies typically underestimate the standing variation present in natural populations and often lack the resolution to pin-point causal variants to a small number of genes. Conversely, association studies are typically underpowered to detect rare alleles, poorly tagged variants, and regions with multiple causal sites in weak LD with one another. MPPs have been proposed to bridge the gap between the above two approaches. Although MPPs have been created in several model systems, only a single MPP has thus far been described in budding yeast. By generating two large, highly outcrossed and genetically heterogeneous populations of *S. cerevisiae* derived from eighteen different founder strains, we have created a powerful resource that can be used in a variety of experimental settings. For instance, these populations can be used in large-scale X-QTL mapping experiments (Ehrenreich *et al*. 2010) to comprehensively dissect the genetic architecture of complex traits as well as large-scale evolve and re-sequence experiments (Lang *et al*. 2011; Parts *et al*. 2011; Burke *et al*. 2014) to determine the mechanisms and course of adaptation to diverse stimuli. Large number of recombinant haploid clones generated from these populations can be used in complementary large-scale I-QTL studies (Bloom *et al*. 2013; Wilkening *et al*. 2014). Due to the high levels of standing variation present, these populations should also prove to be a powerful resource in evolutionary engineering applications, as they are presumably capable of being evolved to carry out a plethora of useful tasks.

The haplotype calling software generated in this study represents a useful resource for the MPP community in general, as it enables highly accurate haplotype calling in poolseq data at reduced coverage. The ability of the algorithm to deal with windows where all founder haplotypes cannot be resolved will have utility in a subset of systems, including our yeast populations. Candidate causal regions can be identified by comparing haplotype frequencies at discrete intervals across the genome in control versus treatment populations. Candidate regions can then be examined in the UCSC genome browser, where genome-wide alignments of all founder strains have been posted. Structural variants can be easily visualized in the browser as can nonsynonymous SNPs, thus pointing investigators to potentially causal genes.

In conclusion, the populations generated in this study represent a novel resource that brings together the power of QTL mapping, the resolution of association studies, and a large amount of natural variation to a model system capable of teasing apart and directly testing the molecular underpinnings of complex traits.

## Acknowledgments

We thank the UC Irvine Genomics High Throughput Facility for the quick turn-around and efficient processing of libraries for sequencing and for help with figuring out the parameters for the Covaris S220 Focused Acoustic Shearer. We would also like to acknowledge our funding source: NIH grant FG18445 to ADL.

## Supplementary Information

**Figure S1.**
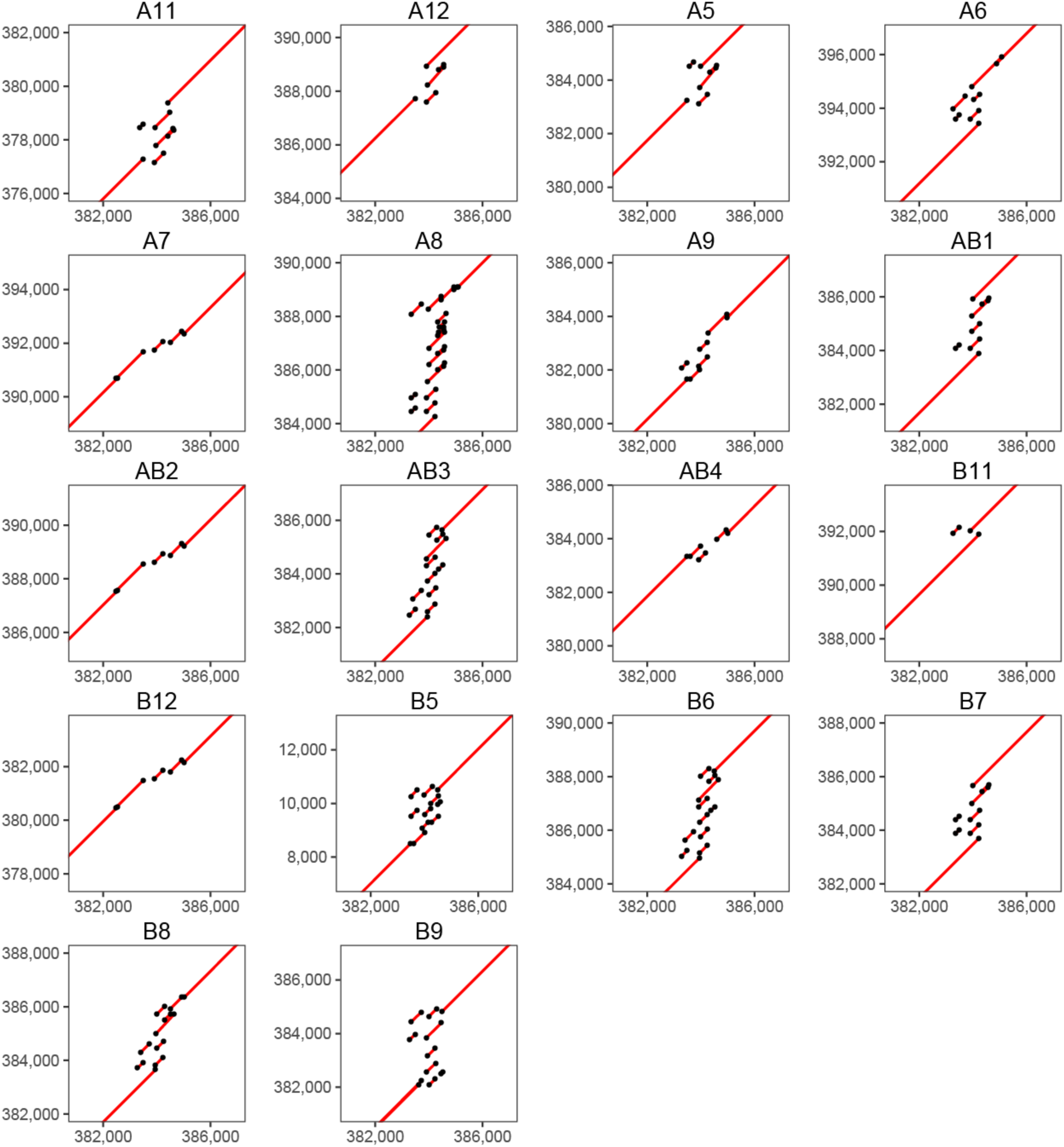
Dot plots showing the numerous structural variants present at this particular site on chromosome IV. Each dot plot was constructed by plotting the sequence alignments of regions from each founder strain (y-axis) against the S288C reference strain (x-axis).

**Figure S2.**
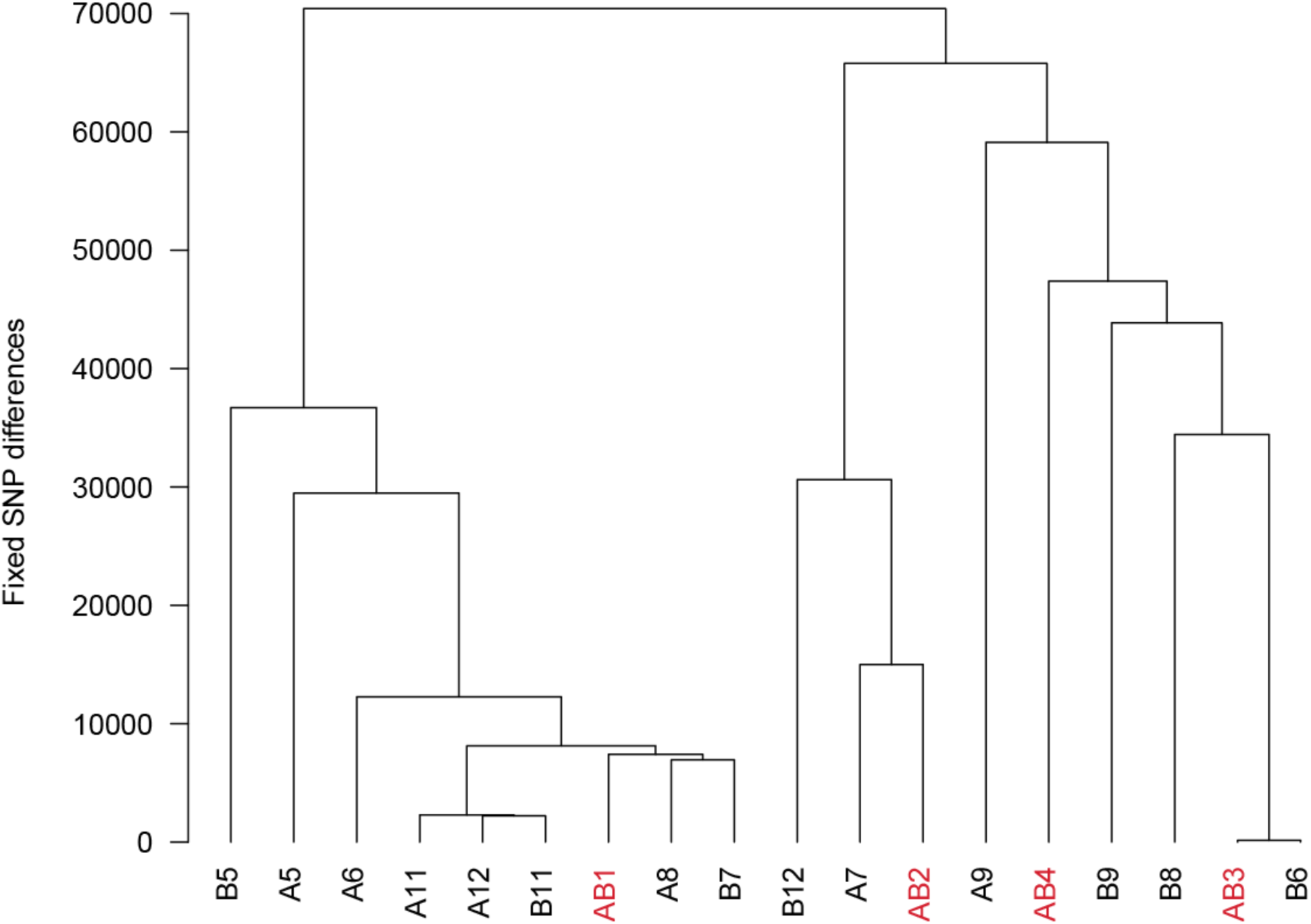
Dendrogram showing the degree of relatedness between the 18 founder strains used in this study. This was constructed in *R* using the *hclust* function on a distance matrix representing the number of differences between each pair of founders. Strains colored in red are the founders from the four-way cross described in Cubillos et al. and used in (Cubillos *et al*. 2013, 2017).

**Figure S3.**
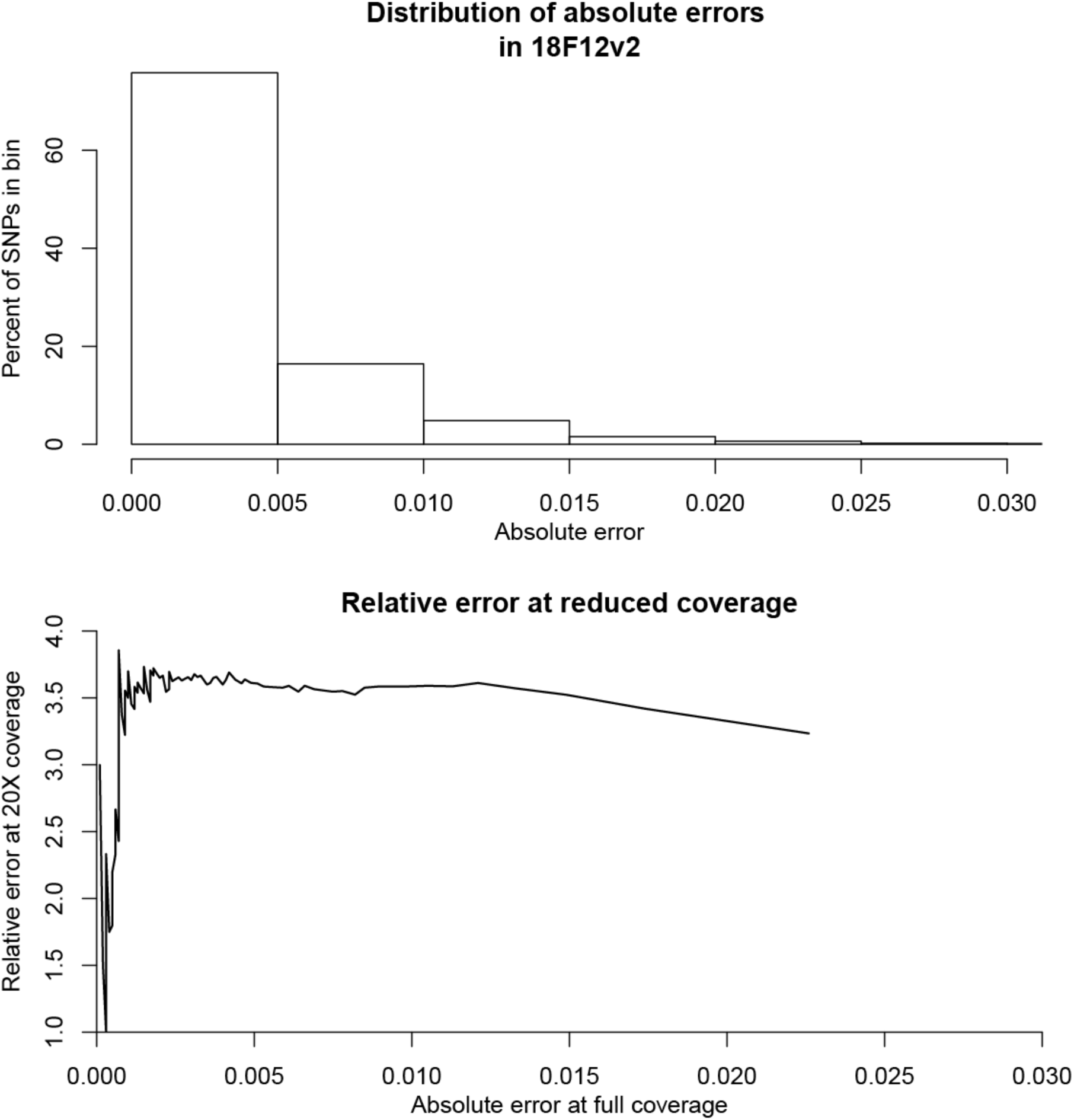
As a measure of the accuracy of the haplotype calling algorithm, we calculated the absolute error between haplotype calls at sites harboring private SNPs and the SNP frequencies in 18F12v2 estimated from 2230X coverage (**A**). The vast majority of sites tested in this manner showed an error rate of less than one percent. We also computed the relative error following downsampling the 18F12v2 data to approximately 20x coverage as a function of the absolute error at full coverage (**B**). At 20X coverage haplotype frequencies are estimated much more accurately than SNP frequencies could be directly estimated at 20X coverage based on binomial sampling.

**Figure S4.**
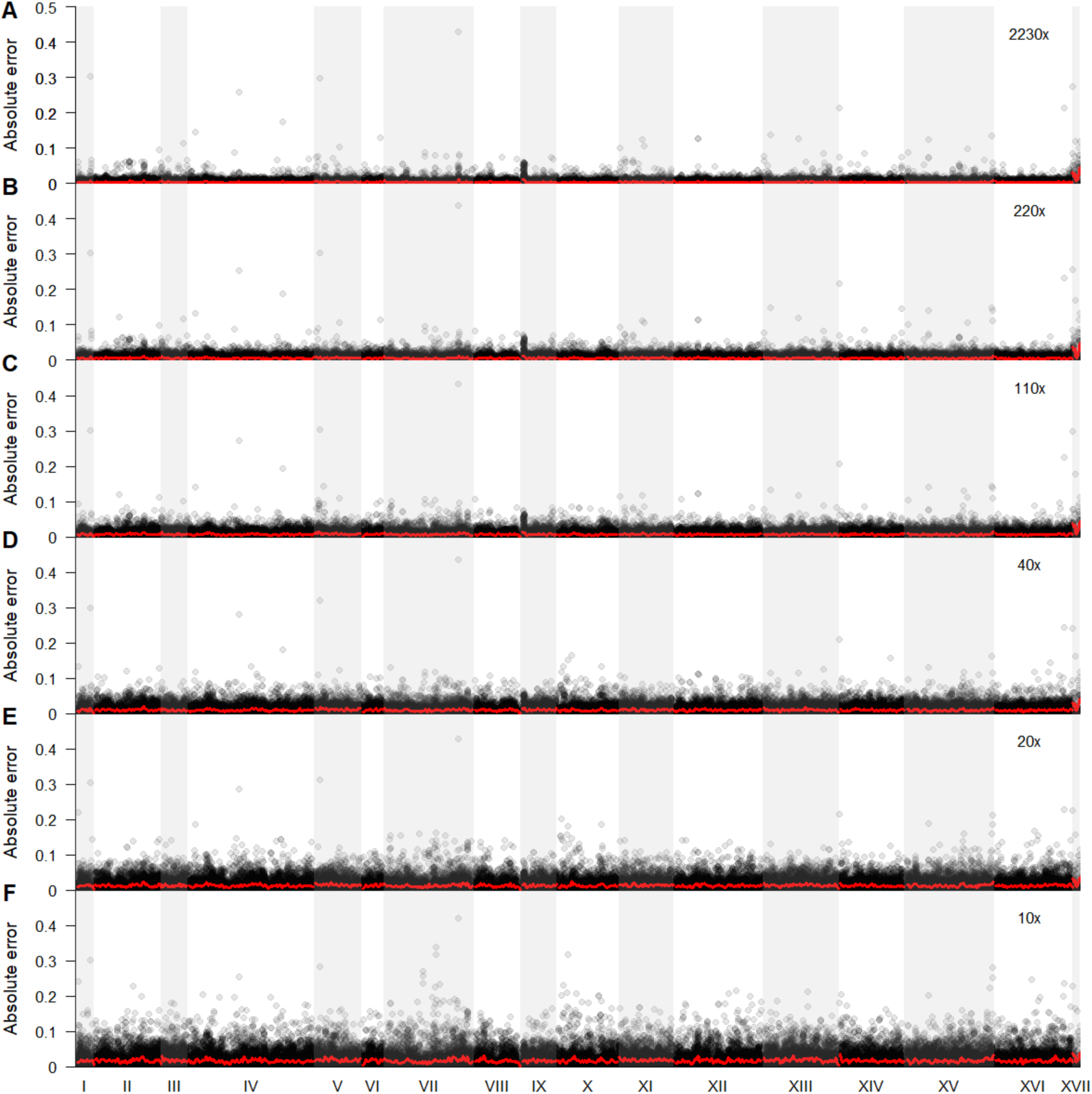
Shown is the absolute error rate between haplotype and SNP frequencies at sites with private SNPs for 18F12v2. Panels A-F represent 100%, 10%, 5%, 2%, 1%, and 0.5% of the full dataset, which approximately corresponds to the coverages shown in the upper right of each panel. The red line represents a kernel regression run using the ksmooth() function in R with kernel set at ‘normal’ and bandwidth set at 20000, approximating the local expected error rate. Despite clear outliers (that occur irrespective of coverage), haplotype frequency estimates are accurate on average.

**Figure S5.**
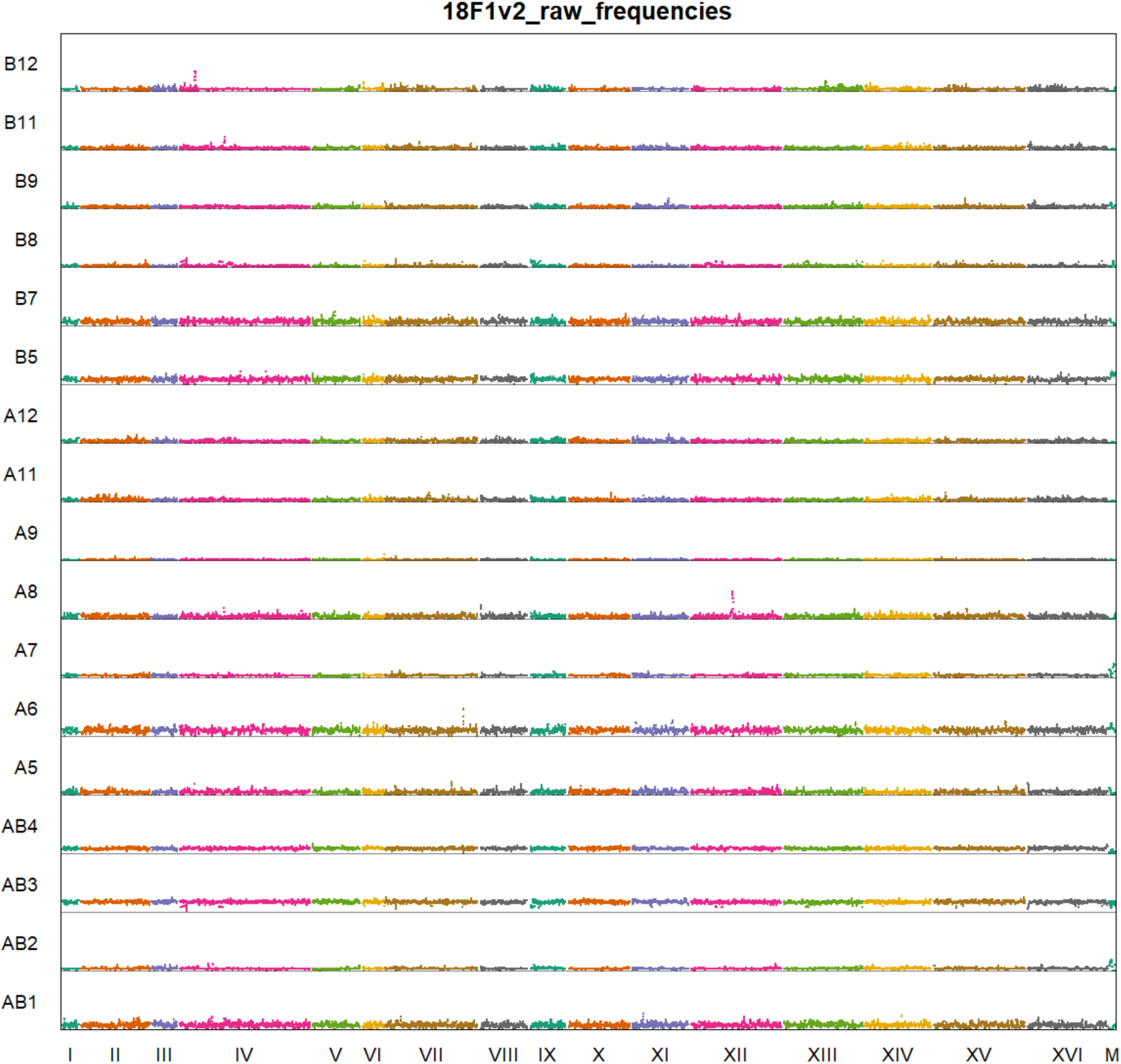
Haplotype frequencies genome-wide for 18F1v2. This sample was taken after the first meiotic generation of version two of the outcrossed population and shows a more balanced distribution of founder haplotypes than the final generation. Haplotype calls for founder B6 were merged with founder AB3, while founders A12 and B11 were merged with founder A11 due their high sequence similarity.

**Figure S6.**
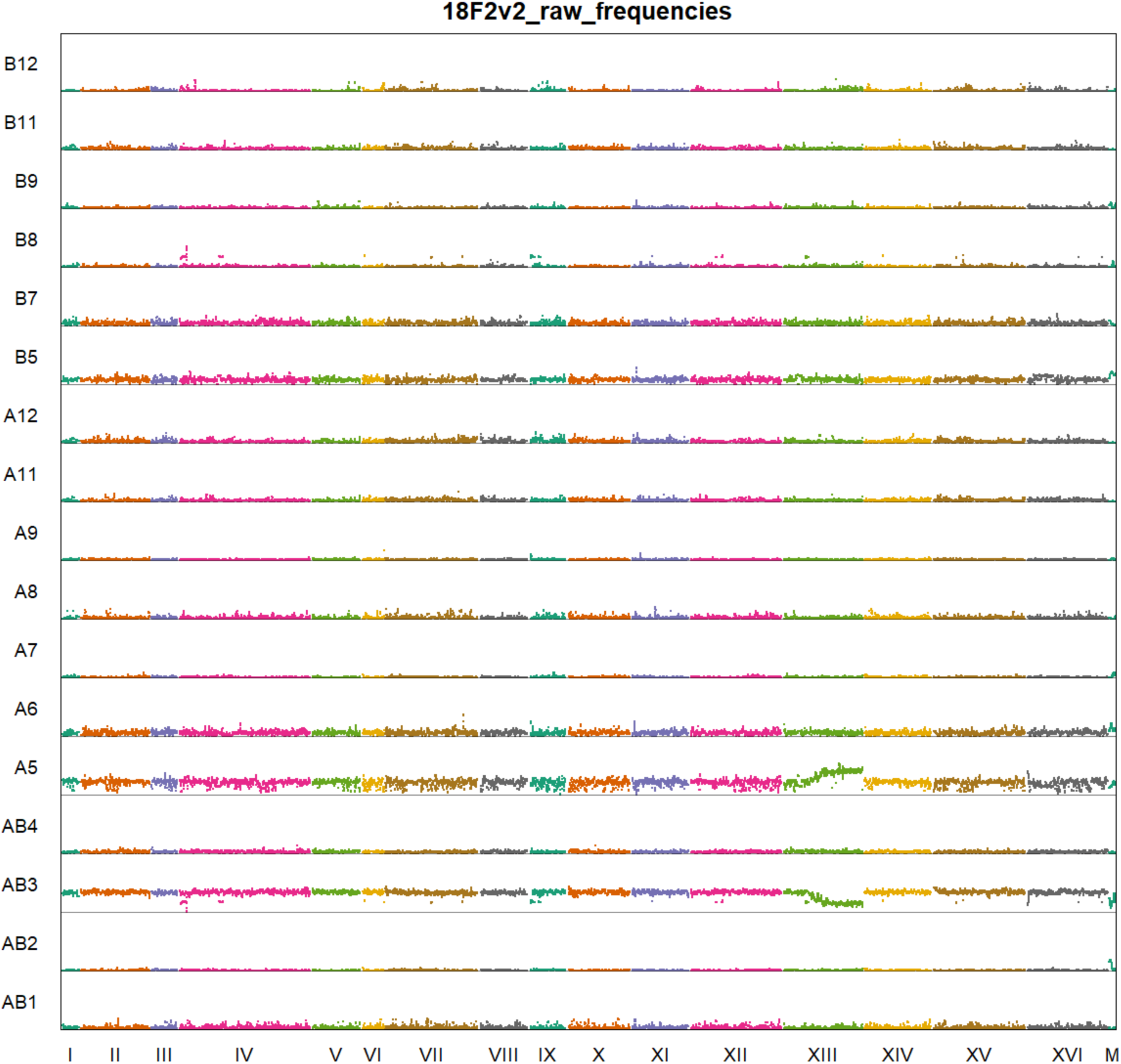
Haplotype frequencies genome-wide for 18F2v2.This sample was taken after the second meiotic generation of version two of the outcrossed population and shows founders AB3 and A5 increasing significantly in frequency as compared to the remaining founders. As above, haplotype calls for founder B6 were merged with founder AB3, while founders A12 and B11 were merged with founder A11.

**Figure S7.**
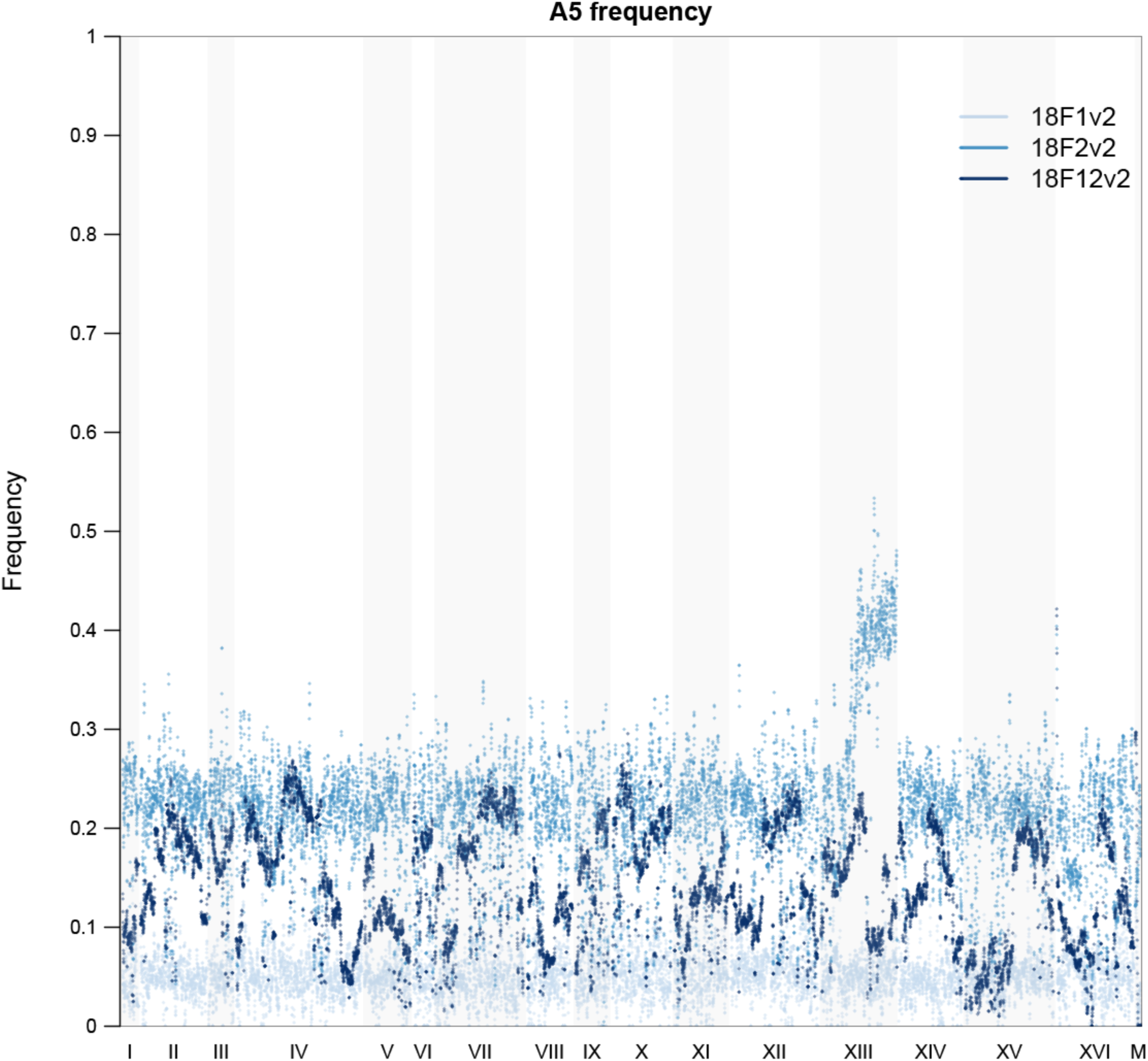
Haplotype frequency genome-wide for the A5 founding strain across the first (18F1v2), second (18F2v2), and twelfth (18F12v2) meiotic generations of the version 2 outcrossed population.

**Figure S8.**
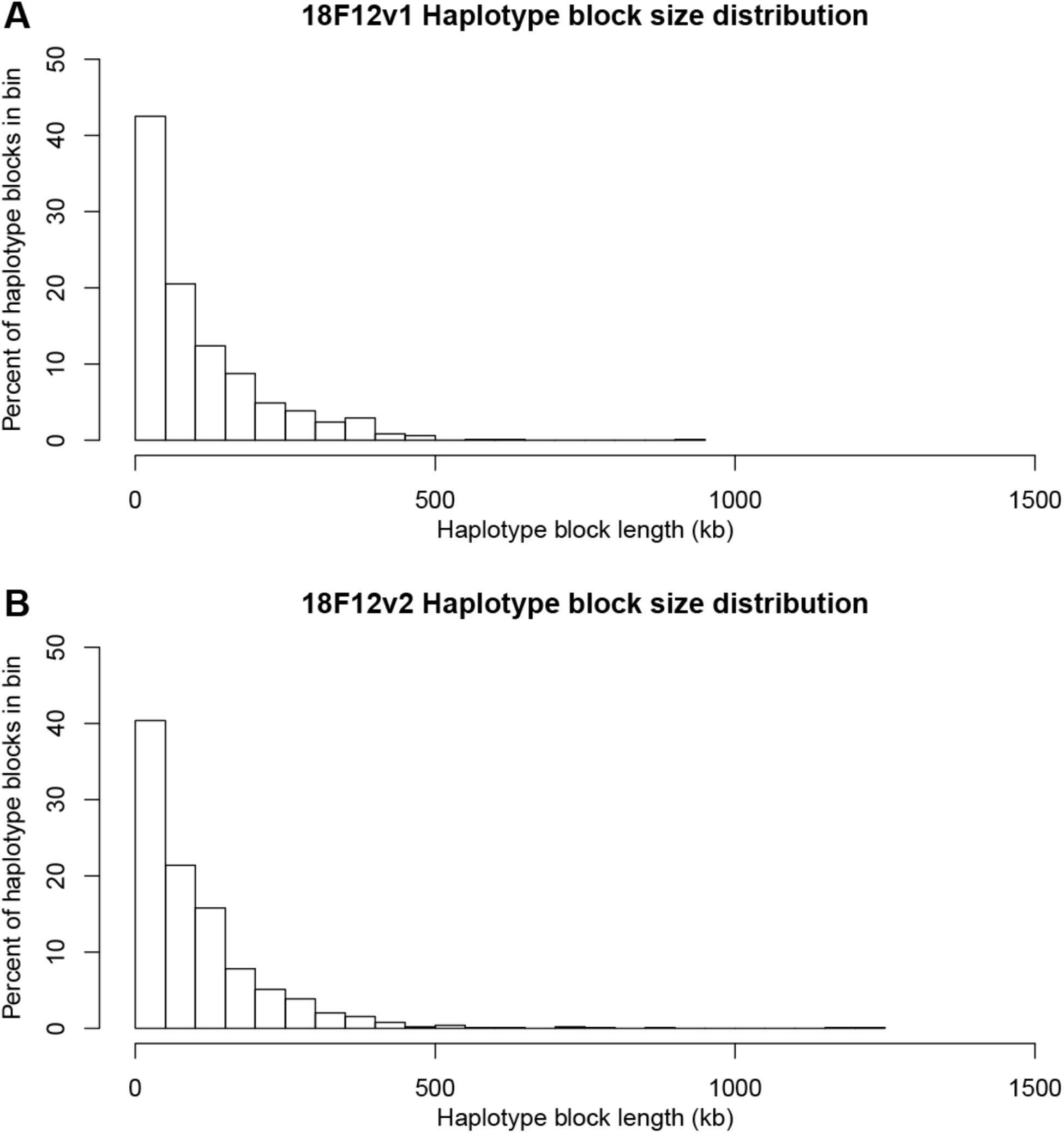
Distribution of haplotype block lengths in recombinant haploid clones derived from 18F12v1 (**A**) and 18F12v2 (**B**). All inferred haplotype blocks, including those called as ties between multiple founding strains, were included in this analysis.

**Figure S9.**
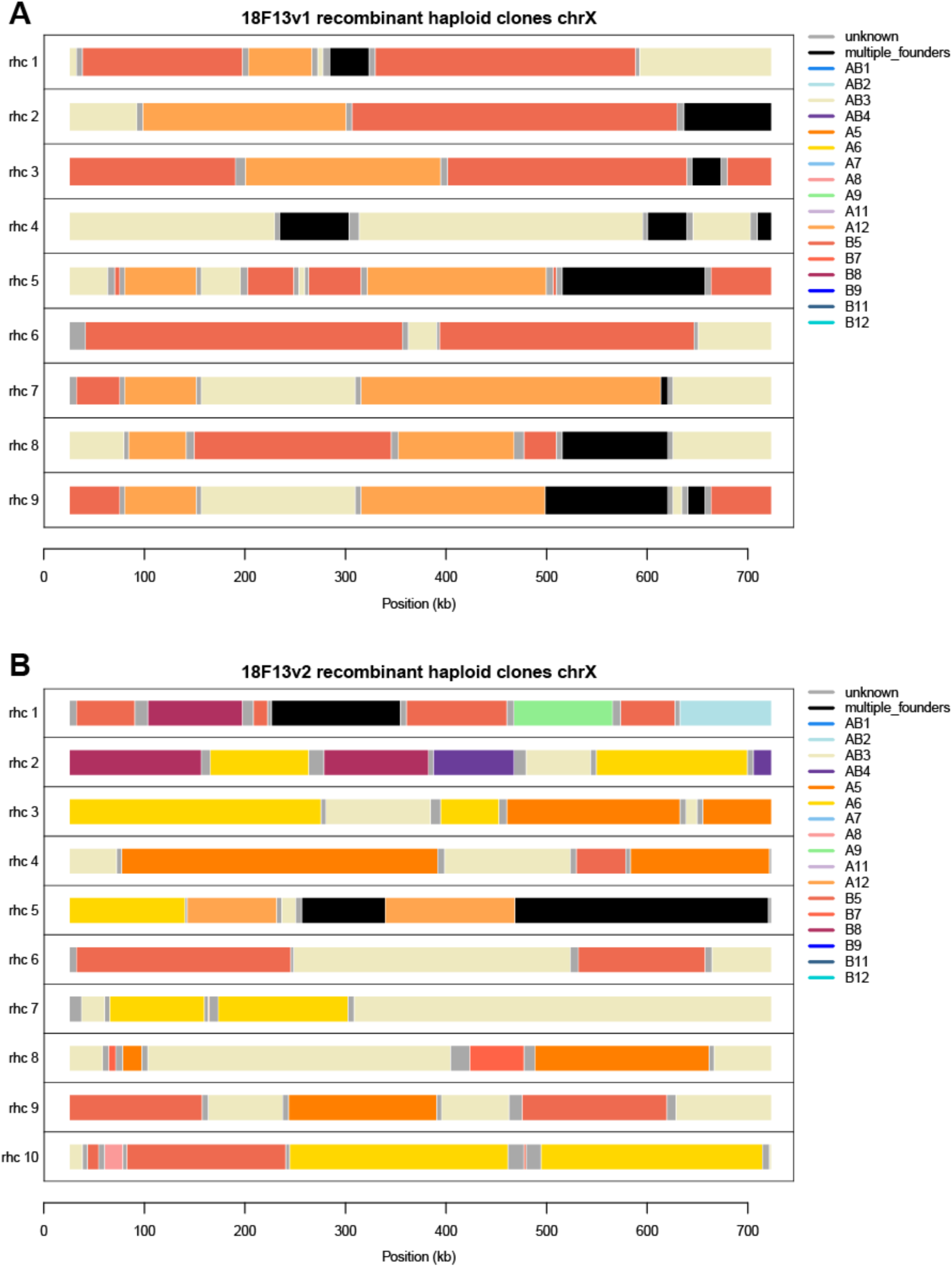
Haplotype diversity at chromosome XII present in 18F13v1 recombinant haploid clones (**A**) and 18F13v2 recombinant haploid clones (**B**).

**Table S1.**
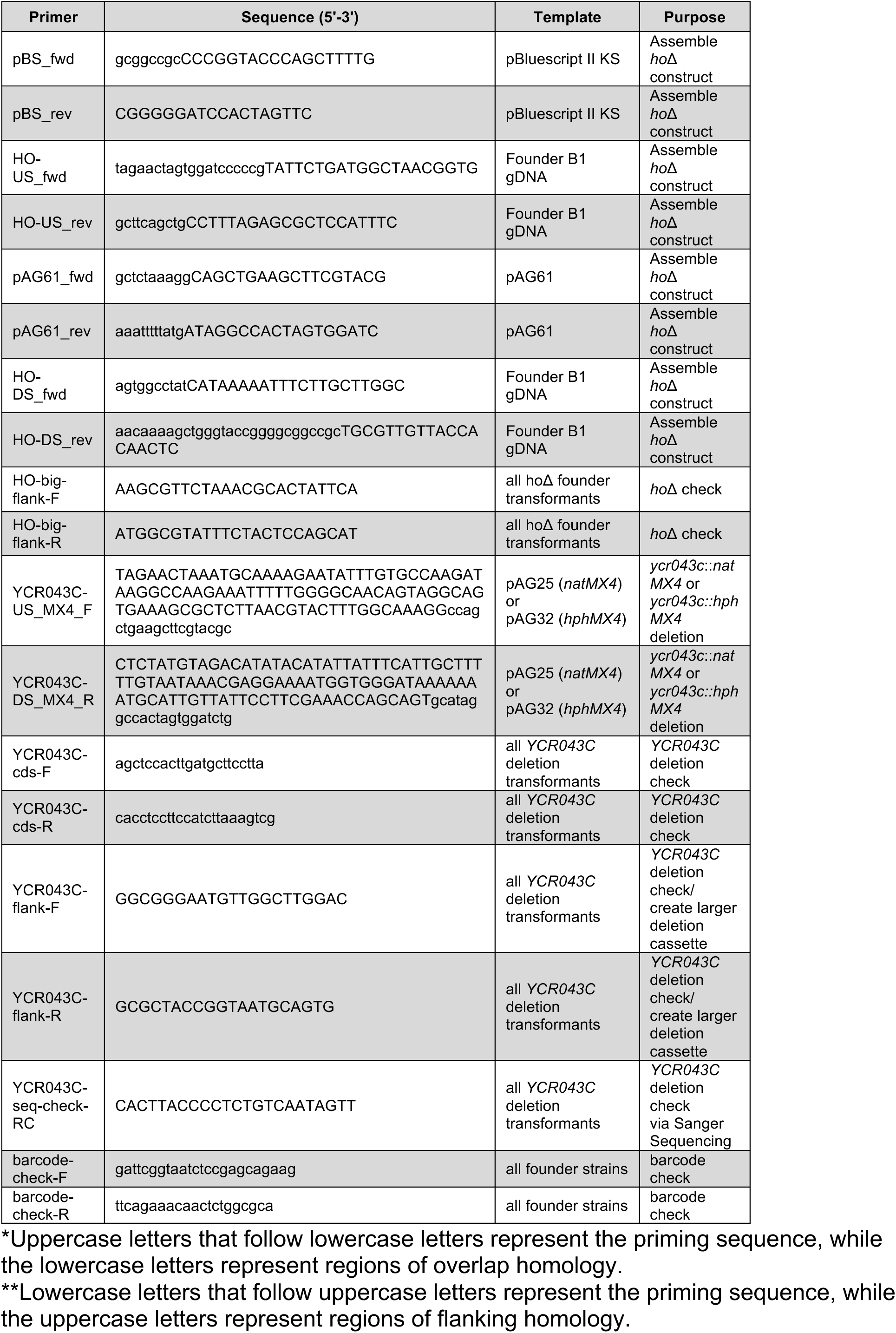
List of primers used in this study.

**Table S2.** Sporulation efficiency of all crosses*

|  | B1 | B2 | B3 | B4 | B5 | B6 | B7 | B8 | B9 | B11 | B12 |
| --- | --- | --- | --- | --- | --- | --- | --- | --- | --- | --- | --- |
| A1 | 0 | 4 | 1 | 3 | 3 | 4 | 3 | 4 | 4 | 1 | 4 |
| A2 | 2 | 1 | 4 | 1 | 4 | 4 | 4 | 3 | 2 | 1 | 3 |
| A3 | 5 | 3 | 3 | ND** | 4 | 3 | 4 | 0 | 2 | 4 | 4 |
| A4 | 4 | 4 | 3 | 4 | 3 | 2 | 4 | 4 | 3 | 3 | 4 |
| A5 | 3 | 2 | 3 | 1 | 1 | 2 | 4 | 2 | 0 | 0 | 1 |
| A6 | 3 | 4 | 4 | 3 | 3 | 4 | 4 | 4 | 3 | 2 | 4 |
| A7 | 3 | 4 | 5 | 4 | 4 | 5 | 5 | 5 | 4 | 4 | 5 |
| A8 | 3 | 3 | 3 | 4 | 3 | 2 | 4 | 0 | 4 | 0 | 4 |
| A9 | 5 | 4 | 3 | 3 | 4 | 2 | 4 | 1 | 2 | 4 | 4 |
| A11 | 4 | 4 | 4 | 4 | 4 | 5 | 4 | 3 | 4 | 1 | 4 |
| A12 | 4 | 4 | 4 | 3 | 3 | 3 | 4 | 4 | 4 | 0 | 4 |

\*A score of 0 means no asci were visible in the field of view while a score of 5 means approximately 90% or more of the objects in the field of view were asci.
\*\*No data

**Table S3.**
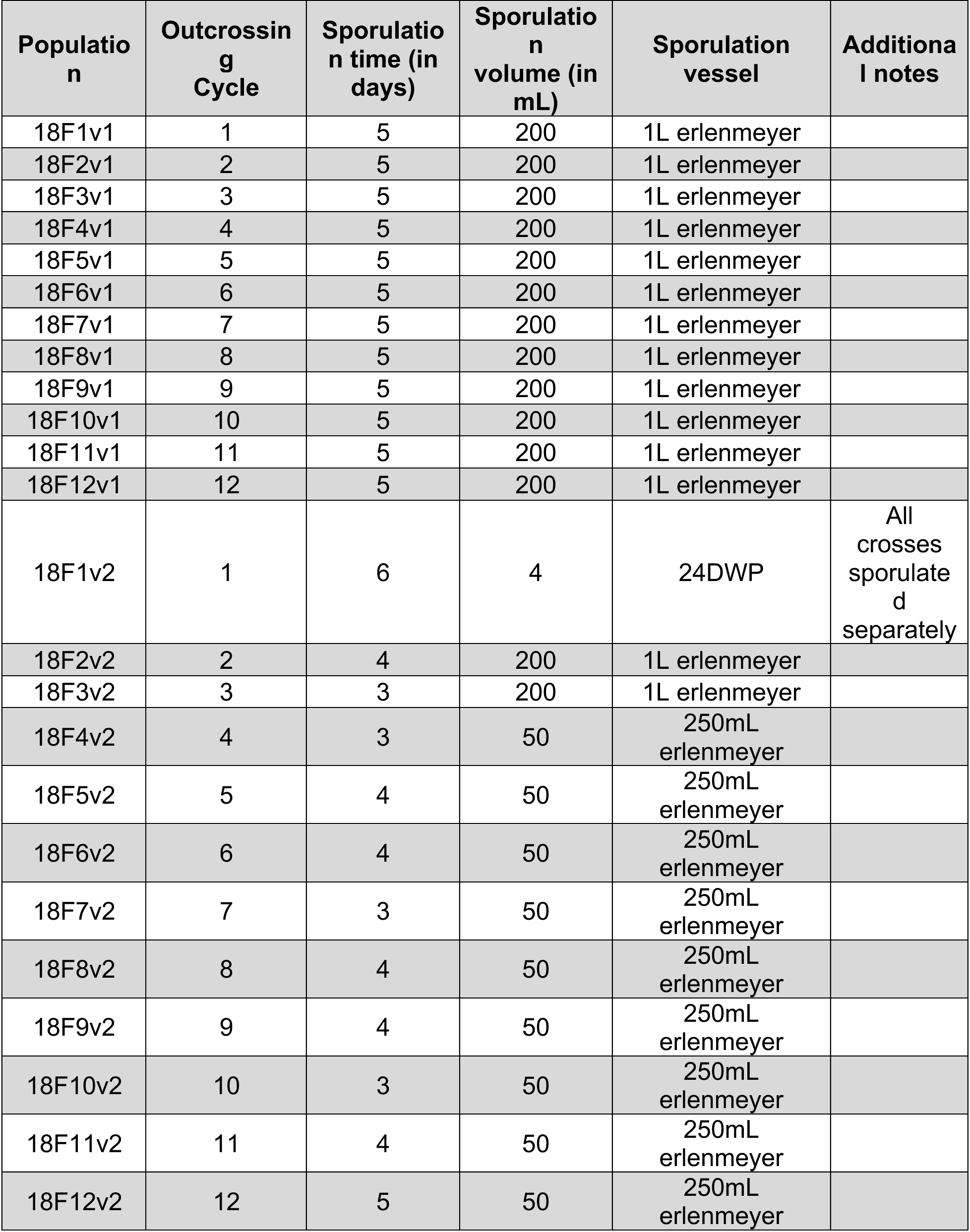
Specific sporulation conditions used in this study.

**Table S4.** All pairwise SNP differences between founder strains

|  | AB1 | AB2 | AB3 | AB4 | A5 | A6 | A7 | A8 | A9 |
| --- | --- | --- | --- | --- | --- | --- | --- | --- | --- |
| AB1 | 0 | 68178 | 60061 | 66679 | 27765 | 12131 | 63233 | 7410 | 70076 |
| AB2 | 68178 | 0 | 48848 | 55578 | 64142 | 67101 | 15008 | 68575 | 63395 |
| AB3 | 60061 | 48848 | 0 | 40134 | 53034 | 58437 | 48360 | 60383 | 52287 |
| AB4 | 66679 | 55578 | 40134 | 0 | 54606 | 64986 | 50777 | 67027 | 59111 |
| A5 | 27765 | 64142 | 53034 | 54606 | 0 | 29485 | 59901 | 27980 | 66127 |
| A6 | 12131 | 67101 | 58437 | 64986 | 29485 | 0 | 62410 | 12281 | 69433 |
| A7 | 63233 | 15008 | 48360 | 50777 | 59901 | 62410 | 0 | 12281 | 63205 |
| A8 | 7410 | 68575 | 60383 | 67027 | 27980 | 12281 | 63629 | 0 | 70409 |
| A9 | 70076 | 63395 | 52287 | 59111 | 66127 | 69433 | 63205 | 70409 | 0 |
| A11 | 8140 | 68612 | 60416 | 67088 | 28220 | 12261 | 63669 | 7898 | 70395 |
| A12 | 7988 | 68533 | 60365 | 67049 | 27943 | 12154 | 63594 | 7794 | 70361 |
| B5 | 28183 | 63545 | 53668 | 55176 | 36714 | 30476 | 59484 | 28362 | 66654 |
| B6 | 60023 | 48876 | 157* | 40176 | 53023 | 58407 | 48373 | 60347 | 52315 |
| B7 | 7320 | 68536 | 60350 | 66977 | 27893 | 12207 | 63582 | 6964 | 70372 |
| B8 | 58073 | 52026 | 34405 | 44614 | 54177 | 56026 | 51878 | 58356 | 53822 |
| B9 | 54032 | 56216 | 43784 | 47394 | 49980 | 53422 | 54434 | 54244 | 57068 |
| B11 | 8083 | 68613 | 60397 | 67067 | 28111 | 12214 | 63672 | 7807 | 70382 |
| B12 | 45831 | 25006 | 52332 | 59231 | 49973 | 46772 | 30619 | 46234 | 65780 |

|  | A11 | A12 | B5 | B6 | B7 | B8 | B9 | B11 | B12 |
| --- | --- | --- | --- | --- | --- | --- | --- | --- | --- |
| AB1 | 8140 | 7988 | 28183 | 60023 | 7320 | 58073 | 54032 | 8083 | 45831 |
| AB2 | 68612 | 68533 | 63545 | 48876 | 68536 | 52026 | 56216 | 68613 | 25006 |
| AB3 | 60416 | 60365 | 53668 | 157* | 60350 | 34405 | 43784 | 60397 | 52332 |
| AB4 | 67088 | 67049 | 55176 | 40176 | 66977 | 44614 | 47394 | 67067 | 59231 |
| A5 | 28220 | 27943 | 36714 | 53023 | 27893 | 54177 | 49980 | 28111 | 49973 |
| A6 | 12261 | 12154 | 30476 | 58407 | 12207 | 56026 | 53422 | 12214 | 46772 |
| A7 | 63669 | 63594 | 59484 | 48373 | 63582 | 51878 | 54434 | 63672 | 30619 |
| A8 | 7898 | 7794 | 28362 | 60347 | 6964 | 58356 | 54244 | 7807 | 46234 |
| A9 | 70395 | 70361 | 66654 | 52315 | 70372 | 53822 | 57068 | 70382 | 65780 |
| A11 | 0 | 2283* | 28464 | 60394 | 7989 | 58473 | 54447 | 2294* | 46503 |
| A12 | 2283* | 0 | 28328 | 60343 | 7885 | 58408 | 54405 | 2217* | 46396 |
| B5 | 28464 | 28328 | 0 | 53655 | 28475 | 54111 | 52160 | 28451 | 49324 |
| B6 | 60394 | 60343 | 53655 | 0 | 60315 | 34439 | 43783 | 60375 | 52328 |
| B7 | 7989 | 7885 | 28475 | 60315 | 0 | 58340 | 54268 | 8006 | 46164 |
| B8 | 58473 | 58408 | 54111 | 34439 | 58340 | 0 | 43869 | 58454 | 54419 |
| B9 | 54447 | 54405 | 52160 | 43783 | 54268 | 43869 | 0 | 54438 | 53416 |
| B11 | 2294* | 2217* | 28451 | 60375 | 8006 | 58454 | 54438 | 0 | 46520 |
| B12 | 46503 | 46396 | 49324 | 52328 | 46164 | 54419 | 53416 | 46520 | 0 |

\*Cells bolded and filled in with blue represent founders that were collapsed due to high sequence similarity (AB3/B6; A11/A12/B11). The table was split for readability.

**Table S5.** Mean haplotype frequencies genome-wide through one, two, and twelve rounds of outcrossing in 18F12v2.

| Founder | 18F1v2_frequency | 18F2v2_frequency | 18F12v2_frequency |
| --- | --- | --- | --- |
| AB1 | 8.0% | 3.2% | 1.2% |
| AB2 | 4.4% | 1.3% | 0.5% |
| AB3 | 17.6% | 33.2% | 41.4% |
| AB4 | 9.0% | 3.3% | 3.3% |
| A5 | 5.0% | 21.8% | 14.0% |
| A6 | 10.5% | 6.4% | 14.3% |
| A7 | 4.2% | 1.2% | 0.4% |
| A8 | 4.3% | 2.2% | 0.7% |
| A9 | 1.1% | 1.4% | 0.5% |
| A11 | 3.5% | 2.8% | 1.2% |
| A12 | 3.6% | 3.4% | 1.7% |
| B5 | 9.1% | 8.4% | 11.3% |
| B7 | 7.2% | 4.6% | 1.0% |
| B8 | 2.3% | 1.4% | 5.7% |
| B9 | 3.2% | 1.3% | 0.8% |
| B11 | 3.1% | 2.6% | 1.3% |
| B12 | 3.8% | 1.4% | 0.6% |

**Note S1.** Strains A9, B1, and B9 all grew slowly after O/N incubation, so all 5mL from each culture was transferred to 5mL of YPD and incubated an additional 3h at 30°C at 250 RPM. During this time, the rest of the haploid strains remained at RT. For mating, 200uL of culture from these slow-growing strains was mixed with 100uL of culture from the rest of the haploid strains..
**Note S2.** Before taking OD630 measurements, the following crosses were accidentally combined: A5xB6 with A5xB7, A6xB2 with A6xB3, A6xB8 with A6xB4, and A11xB7 with A11xB6. To correct for this, twice of much of the combined cultures were added to the final spore pool after normalizing cell density.
**Note S3.** Founder strains A5, A7-A9, A11, A12 and B5-B12 (excluding B10) were sequenced using PE100 reads, while founders A1-A4 and A6 were sequenced using PE150 reads.
**Note S4.** In one of the recombinant haploid clones derived from 18F12v2 (rhc 10), the entirety of chromosome I was called as unknown. Closer examination showed that this chromosome is duplicated and heterozygous for two founding strains, suggesting a partial diploidization event.

## References

Aylor, D. L., W. Valdar, W. Foulds-Mathes, R. J. Buus, R. A. Verdugo et al., 2011 Genetic analysis of complex traits in the emerging Collaborative Cross. Genome Res. 21: 1213–22.

Barghi, N., and C. Schlötterer, 2019 Shifting the paradigm in Evolve and Resequence studies: From analysis of single nucleotide polymorphisms to selected haplotype blocks. Mol. Ecol. 28: 521–524.

Berg, J. J., A. Harpak, N. Sinnott-Armstrong, A. M. Joergensen, H. Mostafavi et al., 2019 Reduced signal for polygenic adaptation of height in UK Biobank. Elife 8:.

Bloom, J. S., J. Boocock, S. Treusch, M. J. Sadhu, L. Day et al., 2019 Rare variants contribute disproportionately to quantitative trait variation in yeast. bioRxiv 607291.

Bloom, J. S., I. M. Ehrenreich, W. Loo, T.-L. V. Lite, and L. Kruglyak, 2013 Finding the sources of missing heritability in a yeast cross. Nature 494: 234.

Bloom, J. S., I. Kotenko, M. J. Sadhu, S. Treusch, F. W. Albert et al., 2015 Genetic interactions contribute less than additive effects to quantitative trait variation in yeast. Nat Commun 6: 8712.

Burke, M. K., G. Liti, and A. D. Long, 2014 Standing genetic variation drives repeatable experimental evolution in outcrossing populations of Saccharomyces cerevisiae. Mol. Biol. Evol. 31: 3228–39.

Chakraborty, M., J. G. Baldwin-Brown, A. D. Long, and J. J. Emerson, 2016 Contiguous and accurate *de novo* assembly of metazoan genomes with modest long read coverage. Nucleic Acids Res. 44: gkw654.

Chakraborty, M., J. J. Emerson, S. J. Macdonald, and A. D. Long, 2019 Structural variants exhibit widespread allelic heterogeneity and shape variation in complex traits. Nat. Commun. 10:.

Chen, G.-B., S. H. Lee, M. R. Robinson, M. Trzaskowski, Z.-X. Zhu et al., 2017 Across-cohort QC analyses of GWAS summary statistics from complex traits. Eur. J. Hum. Genet. 25: 137–146.

Cingolani, P., A. Platts, L. L. Wang, M. Coon, T. Nguyen et al., 2012 A program for annotating and predicting the effects of single nucleotide polymorphisms, SnpEff: SNPs in the genome of Drosophila melanogaster strain w1118; iso-2; iso-3. Fly (Austin). 6: 80–92.

Cubillos, F. A., C. Brice, J. Molinet, S. Tisné, V. Abarca et al., 2017 Identification of Nitrogen Consumption Genetic Variants in Yeast Through QTL Mapping and Bulk Segregant RNA-Seq Analyses. G3 (Bethesda). 7: 1693–1705.

Cubillos, F. A., E. J. Louis, and G. Liti, 2009 Generation of a large set of genetically tractable haploid and diploid *Saccharomyces* â€ƒstrains. FEMS Yeast Res. 9: 1217–1225.

Cubillos, F. A., L. Parts, F. Salinas, A. Bergström, E. Scovacricchi et al., 2013 High-resolution mapping of complex traits with a four-parent advanced intercross yeast population. Genetics 195: 1141–55.

Cullen, P. J., W. Sabbagh, E. Graham, M. M. Irick, E. K. Van Olden et al., 2004 A signaling mucin at the head of the Cdc42- and MAPK-dependent filamentous growth pathway in yeast. Genes Dev. 18: 1695–1708.

Dimitrov, L. N., R. B. Brem, L. Kruglyak, and D. E. Gottschling, 2009 Polymorphisms in multiple genes contribute to the spontaneous mitochondrial genome instability of Saccharomyces cerevisiae S288C strains. Genetics 183: 365–383.

Ehrenreich, I. M., N. Torabi, Y. Jia, J. Kent, S. Martis et al., 2010 Dissection of genetically complex traits with extremely large pools of yeast segregants. Nature 464: 1039–42.

Flint, J., and R. Mott, 2001 Finding the molecular basis of quantitative traits: successes and pitfalls. Nat. Rev. Genet. 2: 437–445.

Garrison, E., and G. Marth, 2012a Haplotype-based variant detection from short-read sequencing.

Garrison, E., and G. Marth, 2012b Haplotype-based variant detection from short-read sequencing.

Gimble, F. S., and J. Thorner, 1992 Homing of a DNA endonuclease gene by meiotic gene conversion in Saccharomyces cerevisiae. Nature 357: 301–306.

Hehir-Kwa, J. Y., T. Marschall, W. P. Kloosterman, L. C. Francioli, J. A. Baaijens et al., 2016 A high-quality human reference panel reveals the complexity and distribution of genomic structural variants. Nat. Commun. 7: 12989.

Huang, X., M.-J. Paulo, M. Boer, S. Effgen, P. Keizer et al., 2011 Analysis of natural allelic variation in Arabidopsis using a multiparent recombinant inbred line population. Proc. Natl. Acad. Sci. U. S. A. 108: 4488.

Kane, P. M., 2007 The long physiological reach of the yeast vacuolar H+-ATPase. J. Bioenerg. Biomembr. 39: 415–421.

Kessner, D., T. L. Turner, and J. Novembre, 2013 Maximum Likelihood Estimation of Frequencies of Known Haplotypes from Pooled Sequence Data. Mol. Biol. Evol. 30: 1145–1158.

King, E. G., S. J. Macdonald, and A. D. Long, 2012a Properties and power of the Drosophila Synthetic Population Resource for the routine dissection of complex traits. Genetics 191: 935–49.

King, E. G., C. M. Merkes, C. L. McNeil, S. R. Hoofer, S. Sen et al., 2012b Genetic dissection of a model complex trait using the Drosophila Synthetic Population Resource. Genome Res. 22: 1558–66.

de Koning, D.-J., and L. M. McIntyre, 2017 Back to the Future: Multiparent Populations Provide the Key to Unlocking the Genetic Basis of Complex Traits. Genetics 206: 527–529.

Koren, S., B. P. Walenz, K. Berlin, J. R. Miller, N. H. Bergman et al., 2017 Canu: scalable and accurate long-read assembly via adaptive k-mer weighting and repeat separation. Genome Res. 27: 722–736.

Kover, P. X., W. Valdar, J. Trakalo, N. Scarcelli, I. M. Ehrenreich et al., 2009 A Multiparent Advanced Generation Inter-Cross to Fine-Map Quantitative Traits in Arabidopsis thaliana (R. Mauricio, Ed.). PLoS Genet. 5: e1000551.

Kowalec, P., M. Grynberg, B. Pajak, A. Socha, K. Winiarska et al., 2015a Newly identified protein Imi1 affects mitochondrial integrity and glutathione homeostasis in Saccharomyces cerevisiae. FEMS Yeast Res. 15:.

Kowalec, P., M. Grynberg, B. Pająk, A. Socha, K. Winiarska et al., 2015b Newly identified protein Imi1 affects mitochondrial integrity and glutathione homeostasis in *Saccharomyces cerevisiae* (I. Dawes, Ed.). FEMS Yeast Res. 15: fov048.

Lander, E. S., and D. Botstein, 1989 Mapping mendelian factors underlying quantitative traits using RFLP linkage maps. Genetics 121: 185–99.

Lang, G. I., D. Botstein, and M. M. Desai, 2011 Genetic variation and the fate of beneficial mutations in asexual populations. Genetics 188: 647–661.

Langmead, B., and S. L. Salzberg, 2012 Fast gapped-read alignment with Bowtie 2. Nat. Methods 9: 357–359.

Li, H., 2011 A statistical framework for SNP calling, mutation discovery, association mapping and population genetical parameter estimation from sequencing data. Bioinformatics 27: 2987–2993.

Li, H., 2013 Aligning sequence reads, clone sequences and assembly contigs with BWA-MEM.

Li, H., and R. Durbin, 2009 Fast and accurate short read alignment with Burrows-Wheeler transform. Bioinformatics 25: 1754–1760.

Li, H., B. Handsaker, A. Wysoker, T. Fennell, J. Ruan et al., 2009 The Sequence Alignment/Map format and SAMtools. Bioinformatics 25: 2078–2079.

Liti, G., and E. J. Louis, 2012 Advances in quantitative trait analysis in yeast. PLoS Genet. 8: e1002912.

Long, Q., D. C. Jeffares, Q. Zhang, K. Ye, V. Nizhynska et al., 2011 PoolHap: Inferring Haplotype Frequencies from Pooled Samples by Next Generation Sequencing (T. Mailund, Ed.). PLoS One 6: e15292.

Long, A. D., S. J. Macdonald, and E. G. King, 2014 Dissecting complex traits using the Drosophila Synthetic Population Resource. Trends Genet. 30: 488–495.

Macdonald, S. J., and A. D. Long, 2007 Joint estimates of quantitative trait locus effect and frequency using synthetic recombinant populations of Drosophila melanogaster. Genetics 176: 1261–81.

Mackay, T. F., 2001 The genetic architecture of quantitative traits. Annu Rev Genet 35: 303–339.

Manolio, T. A., F. S. Collins, N. J. Cox, D. B. Goldstein, L. A. Hindorff et al., 2009 Finding the missing heritability of complex diseases. Nature 461: 747–753.

Marçais, G., A. L. Delcher, A. M. Phillippy, R. Coston, S. L. Salzberg et al., 2018 MUMmer4: A fast and versatile genome alignment system (A. E. Darling, Ed.). PLOS Comput. Biol. 14: e1005944.

Märtens, K., J. Hallin, J. Warringer, G. Liti, and L. Parts, 2016 Predicting quantitative traits from genome and phenome with near perfect accuracy. Nat. Commun. 7: 11512.

McDonald, M. J., D. P. Rice, and M. M. Desai, 2016 Sex speeds adaptation by altering the dynamics of molecular evolution. Nature 531: 233–236.

McMullen, M. D., S. Kresovich, H. S. Villeda, P. Bradbury, H. Li et al., 2009 Genetic properties of the maize nested association mapping population. Science 325: 737–40.

Meersche, K. Van den, K. Soetaert, and D. Van Oevelen, 2009 xsample() : An R Function for Sampling Linear Inverse Problems. J. Stat. Softw. 30:.

Mott, R., C. J. Talbot, M. G. Turri, A. C. Collins, and J. Flint, 2000 A method for fine mapping quantitative trait loci in outbred animal stocks. Proc. Natl. Acad. Sci. U. S. A. 97: 12649–54.

Narasimhan, V., P. Danecek, A. Scally, Y. Xue, C. Tyler-Smith et al., 2016 BCFtools/RoH: A hidden Markov model approach for detecting autozygosity from next-generation sequencing data. Bioinformatics 32: 1749–1751.

Navarro-Aviño, J. P., R. Prasad, V. J. Miralles, R. M. Benito, and R. Serrano, 1999 A proposal for nomenclature of aldehyde dehydrogenases in Saccharomyces cerevisiae and characterization of the stress-inducible ALD2 and ALD3 genes. Yeast 15: 829–842.

Noble, L. M., M. V. Rockman, and H. Teotónio, 2019 Gene-level quantitative trait mapping in an expanded C. elegans multiparent experimental evolution panel. bioRxiv 589432.

O’Rourke, S. M., and I. Herskowitz, 2002 A Third Osmosensing Branch in Saccharomyces cerevisiae Requires the Msb2 Protein and Functions in Parallel with the Sho1 Branch. Mol. Cell. Biol. 22: 4739–4749.

Parts, L., F. A. Cubillos, J. Warringer, K. Jain, F. Salinas et al., 2011 Revealing the genetic structure of a trait by sequencing a population under selection. Genome Res 21: 1131–1138.

Paten, B., M. Diekhans, D. Earl, J. S. John, J. Ma et al., 2011a Cactus graphs for genome comparisons, pp. 469–481 in Journal of Computational Biology,.

Paten, B., D. Earl, N. Nguyen, M. Diekhans, D. Zerbino et al., 2011b Cactus: Algorithms for genome multiple sequence alignment. Genome Res. 21: 1512–1528.

Pritchard, J. K., 2001 Are Rare Variants Responsible for Susceptibility to Complex Diseases? Am. J. Hum. Genet. 69: 124–137.

Ruckenstuhl, C., C. Netzberger, I. Entfellner, D. Carmona-Gutierrez, T. Kickenweiz et al., 2014 Lifespan Extension by Methionine Restriction Requires Autophagy-Dependent Vacuolar Acidification. PLoS Genet. 10:.

Sebastiani, P., N. Solovieff, A. Puca, S. W. Hartley, E. Melista et al., 2011 Retraction. Science 333: 404.

Solares, E. A., M. Chakraborty, D. E. Miller, S. Kalsow, K. Hall et al., 2018 Rapid Low-Cost Assembly of the Drosophila melanogaster Reference Genome Using Low-Coverage, Long-Read Sequencing. G3 (Bethesda). 8: 3143–3154.

Spencer, C. C. A., Z. Su, P. Donnelly, and J. Marchini, 2009 Designing Genome-Wide Association Studies: Sample Size, Power, Imputation, and the Choice of Genotyping Chip (J. D. Storey, Ed.). PLoS Genet. 5: e1000477.

Tan, A., G. R. Abecasis, and H. M. Kang, 2015 Unified representation of genetic variants. Bioinformatics 31: 2202–2204.

The Collaborative Cross, a community resource for the genetic analysis of complex traits, 2004 Nat. Genet. 36: 1133–1137.

Thornton, K. R., A. J. Foran, and A. D. Long, 2013 Properties and Modeling of GWAS when Complex Disease Risk Is Due to Non-Complementing, Deleterious Mutations in Genes of Large Effect (J. K. Pritchard, Ed.). PLoS Genet. 9: e1003258.

Threadgill, D. W., and G. A. Churchill, 2012 Ten years of the Collaborative Cross. Genetics 190: 291–4.

Tkach, J. M., A. Yimit, A. Y. Lee, M. Riffle, M. Costanzo et al., 2012 Dissecting DNA damage response pathways by analysing protein localization and abundance changes during DNA replication stress. Nat. Cell Biol. 14: 966–976.

Visscher, P. M., 2008 Sizing up human height variation. Nat. Genet. 40: 489–490.

Walker, B. J., T. Abeel, T. Shea, M. Priest, A. Abouelliel et al., 2014 Pilon: An Integrated Tool for Comprehensive Microbial Variant Detection and Genome Assembly Improvement (J. Wang, Ed.). PLoS One 9: e112963.

Waterhouse, R. M., M. Seppey, F. A. Simão, M. Manni, P. Ioannidis et al., 2018 BUSCO Applications from Quality Assessments to Gene Prediction and Phylogenomics. Mol. Biol. Evol. 35: 543–548.

Wilkening, S., G. Lin, E. S. Fritsch, M. M. Tekkedil, S. Anders et al., 2014 An evaluation of high-throughput approaches to QTL mapping in Saccharomyces cerevisiae. Genetics 196: 853–865.

WTCCC, 2007 Genome-wide association study of 14,000 cases of seven common diseases and 3,000 shared controls. Nature 447: 661–678.

Ye, C., C. M. Hill, S. Wu, J. Ruan, and Z. Ma, 2016 DBG2OLC: Efficient Assembly of Large Genomes Using Long Erroneous Reads of the Third Generation Sequencing Technologies. Sci. Rep. 6: 31900.

Yue, J.-X., J. Li, L. Aigrain, J. Hallin, K. Persson et al., 2017 Contrasting evolutionary genome dynamics between domesticated and wild yeasts. Nat. Genet. 49: 913–924.

